# NELF coordinates Pol II transcription termination and DNA replication initiation

**DOI:** 10.1101/2024.01.31.578294

**Authors:** Chihiro Nakayama, Yasukazu Daigaku, Yuki Aoi, Qi Fang, Hiroshi Kimura, Ali Shilatifard, Michael Tellier, Takayuki Nojima

## Abstract

Regulation of RNA polymerase II (Pol II) transcription is closely associated with cell proliferation. However, it remains unclear how the Pol II transcription program is altered in cancer to favour cell growth. Here, we find that gene expression of *NELFCD*, a known negative elongation factor, is up-regulated in colorectal tumours. To dissect the direct role of NELF-C on Pol II transcription in such cancer, we employed an auxin-dependent protein degradation system for NELF-C in combination with nascent transcript sequencing technologies. Strikingly, we demonstrated that the acute loss of NELF-C protein globally perturbs Pol II transcription termination and also increases transcription elongation rate, independently of promoter-proximal Pol II pausing. This results in Pol II transcription into DNA replication initiation zones, and may link to failure of the cell cycle transition into S phase. We anticipate that NELF will be a potential therapeutic target to restrict colorectal cancers by promoting transcription-replication conflict.

**HIGHLIGHTS:**

1. Expression of *NELFCD* transcript is up-regulated in colorectal tumors
2. NELF-C protein is mandatory for the transition between G1-S phases during cell cycle
3. NELF-C loss impairs transcription termination independently of Pol II promoter-proximal pausing
4. NELF-C loss leads Pol II to invade DNA replication initiation zones

## INTRODUCTION

Large scale studies of cancer genome sequencing have revealed important somatic mutations that drive cancer evolution. Perturbation of RNA polymerase II (Pol II) transcription by genetic alterations results in changes to the gene expression program that favour tumour proliferation. Notably, dysregulation of Pol II transcription in cancer is associated with transcriptional “addiction”, a process whereby tumours select specific transcription factors to promote cancer cell growth^1^. This dependence results in a higher sensitivity of a cancer cell to the perturbation of specific transcription factors that could be exploited therapeutically. However, it remains largely unclear how Pol II transcriptional programs differ in tumours as compared to normal tissues.

Multiple steps such as initiation, elongation, and termination contribute to the eukaryotic Pol II transcription cycle^1^. After transcription initiation, Pol II is paused at a promoter-proximal region located at 20-40 nucleotides (nt) downstream of the transcription start site (TSS)^2–4^. This step is mainly regulated by two protein complexes; negative elongation factor (NELF) consisting of subunits NELF-A, -B, -C/D, and E, and 5,6-dichloro-1-b-D-ribofuranosylbenzimidazole (DRB)-sensitivity inducing factor (DSIF) comprising SPT4 and SPT5^5,6^. In vitro transcription assay systems have demonstrated that Pol II transcription elongation is suppressed by adding solely NELF or combination of NELF and DSIF proteins^7^. Pol II is then released from such pausing by the cyclin-dependent kinase 9 (CDK9)-mediated phosphorylation of NELF and SPT5^8^. The mechanism of paused Pol II-NELF-DSIF complex was somewhat clarified by recent cryo-electron microscopy (cryo-EM) analysis^9^. An acute depletion of NELF-C protein results in a shift downstream of Pol II pausing to the +1 nucleosome dyad-associated downstream region of the promoter-proximal region^10^. Also, rapid depletion of SPT5 protein causes Pol II degradation, demonstrating that SPT5 contributes to to Pol II stabilization^11^. These findings indicate that NELF and DSIF control Pol II transcription at both pausing and and elongation steps^6^. However, it remains unexplored whether NELF and DSIF also play a role in transcription termination.

At the transcript end site (TES), the endonuclease CPSF73, as a part of the cleavage and polyadenylation (CPA) complex cleaves the nascent RNA at 20-30nt downstream of the polyadenylation signal (PAS, AAUAAA)^12,13^. This CPA RNA cleavage is essential to recruit the nuclear 5’-3’exonuclease, Xrn2 to the 5’end of the downstream cleaved RNA. Xrn2 degrades nascent RNA produced by the elongating Pol II complex to trigger Pol II removal from chromatin. In addition, CPA RNA cleavage is coupled to mRNA polyadenylation that adds a poly(A) tail to the cleaved upstream RNA. This poly(A) tailed RNA is then released from chromatin into the nucleoplasm. The coupling between RNA cleavage and degradation is critical to transcription termination of protein coding (pc) genes^14,15^. On the other hand, Pol II transcription of replication-dependent histone (RDH) genes is terminated independently of PAS and Xrn2. Although CPA factors (such as CPSF73, CPSF100, and CstF64) together with stem-loop binding factor and U7 small nuclear RNA (snRNA) cleave downstream of the RDH transcript, no poly(A) tail is added to its 3’end^12,16^. Another critical RNA cleavage machinery, referred to as the Integrator complex contains IntS11, a homologue of CPSF73. IntS11 cleaves the Pol II nascent transcripts in both small and long noncoding RNAs (lncRNAs) as well as 5’ end regions of pre-mRNAs^17–21^.

Alternative cleavage and polyadenylation (APA) produces multiple mRNA isoforms at the 3’ends of 70% of mammalian pc genes to fine-tune their gene expression^22,23^. In particular APA regulates 3’UTR length, resulting in altered mRNA stability, translation, and localization^24,25^. Several trans-acting proteins have been identified as APA factors. Especially, CstF64^26,27^, Fip1^28^, and PCF11^29^ contribute to efficient RNA cleavage at proximal PAS. Shorter 3’UTR isoform is generally associated with cancer cell proliferation as proximal PAS usage contributes to an increase in oncogenic protein levels in cancer cells due to a decreased microRNA-mediated translational suppression^30^. Pol II transcription speed also contributes to APA^31^. Thus, a slow Pol II mutant tends to select proximal PAS ^32–35^, whereas a fast Pol II mutant switches APA to distal PAS^34^.

To dissect mechanisms of Pol II transcription and its coupled RNA processing, several technologies have been developed to detect newly transcribed RNAs (nascent RNAs)^36^. For example, high throughput sequencing methods for newly synthesized RNAs labelled with modified nucleotides in isolated nucleus, such as precise run-on sequencing (PRO-seq)^2^, showed transcription activity and Pol II pausing. Metabolic RNA labelling with 4-thio Uridine in living cells is also employed to monitor RNA synthesis, called transient transcriptome-sequencing (TT-seq)^37^. Furthermore, we have previously developed the polymerase-intact nascent transcript-sequencing (POINT-seq)^14^ method that profiles TSSs, co-transcriptional RNA cleavage and degradation, co-transcriptional RNA splicing, and read-through transcripts.

Perturbation of transcription termination leads to genomic stresses such as transcription-replication (T-R) conflict principally caused by collision between transcription and replication machineries or between co-transcriptional RNA-DNA hybrid and the incoming replication forks^38^. An siRNA screen of T-R conflict-suppressing factors in human cells identified the CPA factors such as WDR33 that directly binds to the PAS^39^. Notably our previous study showed that siRNA knockdown of the histone chaperone protein SPT6 causes T-R conflict via a genome-wide activation of lncRNA transcription and an impairment of their transcription termination due to a failure of the Integrator complex recruitment to elongating Pol II^40^.

In addition to the effect on the replication fork via T-R conflict, a relationship between replication initiation and transcription has been highlighted by multiple studies^41–43^. Particularly, DNA replication initiation preferentially occurs at genomics regions that neighbour the TSS and TES of actively transcribing genes. The regulation of replication initiation is divided into multiple steps^44^. First, the origin recognition complex (ORC) binds DNA at the replication initiation site and subsequently the pre-loaded replicative helicase Mcm2-7 which assembles with Cdc6 and Cdt1 are recruited to the ORC-bound DNA site. After a double hexamer of Mcm2-7 complexes are topologically loaded onto DNA, the activity of cyclin-dependent kinases (CDKs) and Dbf4-dependent kinases (DDKs) promote formation of the preinitiation complex which includes some components of the replisome GINS, Cdc45, and DNA Polymerase as well as TOPBP1 (Dpb11in yeast). Finally, DNA unwinding is initiated, facilitating formation of the replisome.

In this study we demonstrate that gene expression of NELF-C is significantly up-regulated in colorectal tumours. To investigate the direct role of NELF-C protein in colorectal cancer, we employed an auxin-degron system to acutely degrade NELF-C protein in the human colorectal cancer cell line DLD-1. We observe that loss of NELF-C results in a reversible G1 cell cycle arrest. Strikingly, POINT-seq and TT-seq analysis reveals that acute depletion of NELF-C protein induces an increased transcription elongation rate and a transcription termination defect in pc genes. Notably, loss of NELF-C protein generated extended readthrough transcripts, resulting in Pol II invasion into DNA replication initiation zones. Furthermore, the DNA replication licensing factor CDT1 globally failed to be recruited to chromatin, resulting in an aberrant cell cycle G1-S transition. Overall, our findings provide insight into NELF function at gene 3’ends. We propose that the NELF complex restricts deleterious transcription readthrough to avoid interference of DNA replication firing by dysregulated Pol II.

## RESULTS

### Expression of *NELFCD* transcripts is highly up-regulated in colorectal cancer cells

Colorectal tumours are one of the most frequent cancers types worldwide and associated with patient-specific transcriptional heterogeneity^45,46^. In order to understand transcriptional addiction in colorectal cancer, we re-analyzed RNA-seq data from normal (N) and tumour (T) tissue from the large intestine in Genotype-Tissue Expression (GTEx) and The Cancer Genome Atlas (TCGA) databases (Figure 1A and Supplementary Table 1). For this comparative analysis, we focus on a gene category “Pol II transcription cycle” (104 genes) which consists of initiation (I), elongation (E), and termination (T) (Figure 1B). We found that expression of the *NELFCD* gene transcript, as well as CPA genes such as *CSTF1*, *CSTF2*, and *CPSF3*, is significantly up-regulated in colorectal carcinoma (COAD) and rectum adenocarcinoma (READ), although *SUPT4H1* gene (coding SPT4 protein) is unchanged (Figures 1B-D). Inversely, known inhibitor genes of the cell cycle, such as *CDKN1A* (coding P21 protein) and *CDKN1C* (coding P57 protein), are down-regulated in COAD and other cancer types (Figure 1C and Figure S1A and S1B), consistent with an increased cellular proliferation. While the most significant up-regulation of *NELFCD* occur in COAD and READ, we also observed a slight increase of *NELFCD* expression in other cancer types (Figures 1E and S1B). Similarly, we re-analyzed comparative proteomic data of paired human colon cancer and adjacent normal tissues^47^, detecting upregulated protein levels of all NELF subunits, but not SPT4 protein in primary colon cancer (Figure S1C). This result suggests that the Pol II transcription program in READ and COAD is highly dependent on NELF-CD protein (Figure 1F). Consequently, we decided to investigate NELF-CD function on the Pol II transcription cycle in colorectal cancer.

**Figure 1.**
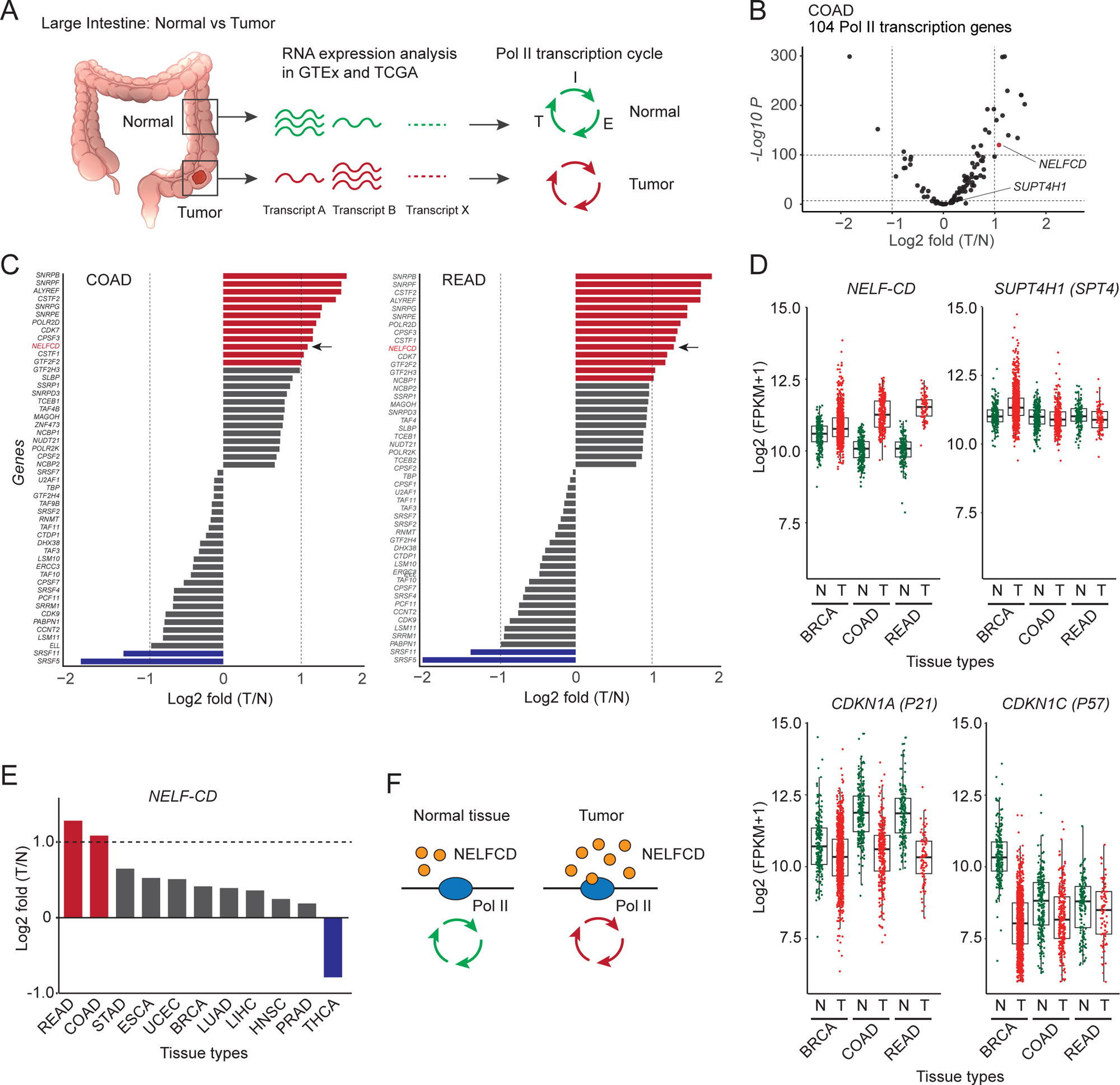
Expression of *NELFCD* transcripts is highly up-regulated in colorectal cancer cells. (A) Schematic diagram of Pol II transcription cycle in Normal (N) tissue and Tumor (T) of large intestine. RNA expression level of Pol II transcription cycle-associated genes in GTEx and TCGA databases were analysed. I: Initiation, E: Elongation, T: Termination. (B) Volcano plot for fold change of T vs N of 104 Pol II transcription-associated genes in COAD. *NELFCD* and *SUPT4H1* genes are indicated with an arrow. *NELFCD* gene is highlighted in red. (C) Log_2_ fold change of T vs N of the Pol II transcription-associated genes. Up-regulated and down-regulated genes are highlighted in red and blue, respectively. *NELFCD* gene is indicated with an arrow. (D) Comparison of RNA expression levels of *NELFCD*, *SUPT4H1*, *CDKN1A*, and *CDKN1C* genes in BRCA (N: n=199, T: n=982), COAD (N: n=153, T: n=87), and READ (N: n=235, T: n=285). (E) Log_2_ fold change of T vs N of *NELFCD* across indicated tissue types. (F) NELF transcription addiction in tumor.

### Nuclear NELF complex controls the cell cycle

NELF-C and NELF-D (the nine amino acids shorter version of NELF-C) are transcriptional isoforms with a different N-terminus domain generated from the *NELFCD* gene. To dissect the function of NELF-C/D, we employed the previously established DLD-1 colorectal cancer cells expressing NELF-C or SPT4 proteins that have been tagged with an auxin-inducible degron (AID) on their C-terminus^10,11^. Notably SPT4 is used as a negative control since it is not up-regulated in colorectal tumors (Figures 1C and S1B). Following addition of auxin (IAA), efficient depletion of NELF-C/D-AID protein (named NELF-C-AID in this study) was detected by western blotting of whole cell extract (WCE) (Figure S2A). Similarly, SPT4-AID protein was specifically degraded upon IAA treatment (Figure S2B). While the degradation of NELF-C protein occurs in less than two hours (2h), other components of the NELF complex are not affected to the exception of NELF-A (Figure S2A), which is known to be closely associated with NELF-C^9^. However, western blot performed on the nuclear fraction reveals that the depletion of NELF-C results in a rapid nuclear loss of NELF-B and NELF-E and at a slower rate of NELF-A (Figure 2A), suggesting that NELF-C stabilizes the nuclear NELF complex. Importantly, the loss of whole NELF complex did not affect the protein and phosphorylation levels of Pol II largest subunit in nuclear fraction (Figure 2A).

**Figure 2.**
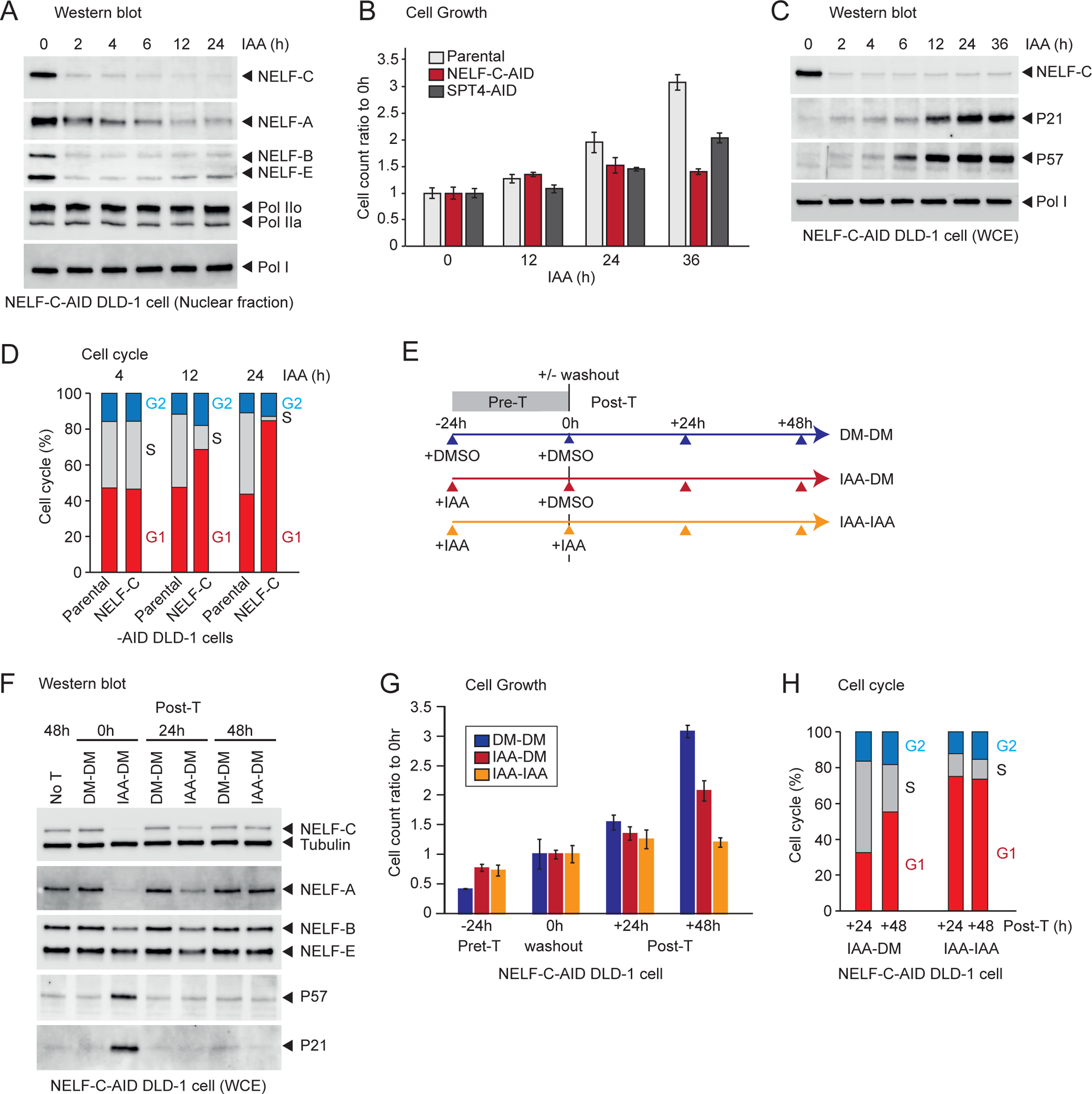
Nuclear NELF complex controls the cell cycle. (A) Western blot of NELF-C-AID DLD-1 cell nuclear fraction using the indicated antibodies. The whole NELF complex is lost from the nucleus following IAA addition. Treatment time (h) of IAA is indicated. (B) Cell count ratios of 12, 24, and 36h IAA to 0h IAA are shown. Parental, NELF-C-AID, and SPT4-AID DLD-1 cells are displayed in light grey, red, and dark grey, respectively. (C) Western blot of NELF-C-AID DLD-1 whole cell extract (WCE) using the indicated antibodies. Treatment time (h) of IAA is indicated. (D) Cell cycle (%) of parental and NELF-C-AID DLD-1 cells following the indicated treatment time (h) of IAA. (E) Schematic timeline of pre-treatment (Pre-T, 24h) and post-treatment (Post-T, up to 48h). For washout, DMSO (DM) or IAA were used. (F) Western blot of NELF-C-AID DLD-1 WCE using the indicated antibodies. No treatment (No-T) and post-treatment time (h) of IAA is indicated. (G) Cell count ratios of -24 (Pre-T), +24 (Post-T), and +48h (Post-T) DMSO or IAA to 0h (washout) are shown. DM-DM, IAA-DM, and IAA-IAA are displayed in blue, red, and orange, respectively. (H) Cell cycle (%) of NELF-C-AID DLD-1 cells (Pre-T, 24h) in the indicated Post-T time (h) of DM or IAA.

Long-term degradation of NELF-C protein has been previously found to result in a severe cell growth defect^10^. However, it remains unclear whether this is due to cell death or cell cycle arrest. To investigate this further, we performed cell growth assays for 36h in the absence or presence of IAA in the NELF-C-AID and SPT4-AID cell lines (Figure 2B). Following IAA treatment, NELF-C-AID cells stopped growing with no detection of dead cells after 12h (Figure 2B, red bars) while SPT4-AID cells continued to proliferate, but slower than parental cells (Figure 2B, grey bars).

To investigate the mechanism behind the growth arrest following NELF-C degradation, we performed western blot of WCE against the known cell cycle inhibitors P21 and P57 (Figure 2C) and observed a clear increase in the expression of these two proteins after 6-12h of IAA treatment. NELF depletion generally decreases gene expression with the exception of fly cells^5,48,49^. Our sequencing analysis of nuclear RNAs that are enriched by oligo dT25 beads (nuclear pA+RNA-seq) following IAA treatment for 4h, 12h, and 24h displayed both up-regulated (4h: 590 genes, 12h: 373 genes, 24h: 404 genes) and down-regulated (4h: 334 genes, 12h: 301 genes, 24h: 257 genes) gene expression in human DLD-1 cells (Figure S2C). The massive increase in expression of the cell cycle inhibitors *CDKN1A* (P21), *CDKN1C* (P57), and *CDKN2B* (P15) transcripts was confirmed by nuclear pA+RNA-seq in the absence of NELF-C protein. In contrast RNA expression of the housekeeping *GAPDH* gene was unchanged (Figure S2D). Concomitantly with expression of cell cycle inhibitors and growth arrest after 6-12h of IAA treatment, we observed an increase in nuclear γH2AX, a DNA damage marker protein (Figure S2E).

These observations led us to investigate the effect of NELF-C protein degradation on the cell cycle. Strikingly, our cell sorting analysis displayed a large increase in G1 phase coupled with a concomitant decrease in S phase cell populations following 12h IAA treatment (Figure 2D). To investigate whether the effect of NELF-C loss on cell growth and cell cycle is reversible, we performed washout experiments where following 24h of pre-treatment with IAA to degrade NELF-C, IAA was replaced by DMSO (washout) and the cells then monitored for 48h in post-treatment (Figure 2E). By western blot, we observed a partial and then total recovery of NELF-C and NELF-A protein levels at 24h and 48h, respectively (Figure 2F). Simultaneously to the recovery of NELF protein levels, we also observed a complete reduction to background levels of P21 and P57 proteins from 24h post-treatment. This loss of the cell cycle inhibitors in the washout condition is also associated with a partial recovery in the growth rate after 48h (Figure 2G) and a complete recovery of the proportion of cells in G1 and S phases at 24h in post-treatment (Figure 2H). Together, our data indicate that the nuclear NELF complex is required for establishment of an efficient cell cycle progression.

### Acute NELF depletion fails to terminate Pol II transcription

In order to dissect the function of NELF and DSIF complexes on elongating Pol II in colorectal cancer cells, we employed POINT-seq technology in NELF-C-AID, NELF-E-AID, and SPT4-AID DLD-1 cells. POINT analysis involves the isolation of nascent Pol II transcripts by immunoprecipitation with Pol II CTD antibody from a DNase-digested chromatin fraction, highly purified using 1M Urea and 3% Empigen detergent (Figure 3A). Our POINT-seq method detected a substantial transcription readthrough of pc gene, *RPS23* in control DMSO-treated DLD-1 cells (Figure 3B). Importantly, 4h IAA treatment in NELF-C-AID cells significantly extended such transcription readthrough, but not in NELF-E-AID cells. The 4h depletion of NELF-E protein partially (∼50%) reduced the core NELF-B and NELF-C protein levels in the nuclear fraction. This may result in less impact of NELF-E protein on Pol II transcription termination (Figure S3A). On the other hand, 3h SPT4 depletion decreases POINT-seq signals throughout the transcription unit, indicating that SPT4 enhances transcription elongation. These transcriptional effects are detected genome-wide by metagene analysis of nonoverlapping pc genes (Figure 3C, n=17,963). To determine the extent of the transcription termination defect, we quantified a ratio of POINT-seq signals in the 2.5kb downstream region of PAS over the gene body (TSS to TES) to determine a termination index (TI). In two biological replicates, our POINT-seq analysis detected higher TI in 4h IAA than in 4h DMSO treatments in NELF-C-AID DLD-1 cells, but not in the NELF-E-AID and SPT4-AID cells (Figure 3D). This result indicates that the NELF core complex (minus NELF-E) enhances Pol II termination on a subset of pc genes, independently of the DSIF complex.

**Figure 3.**
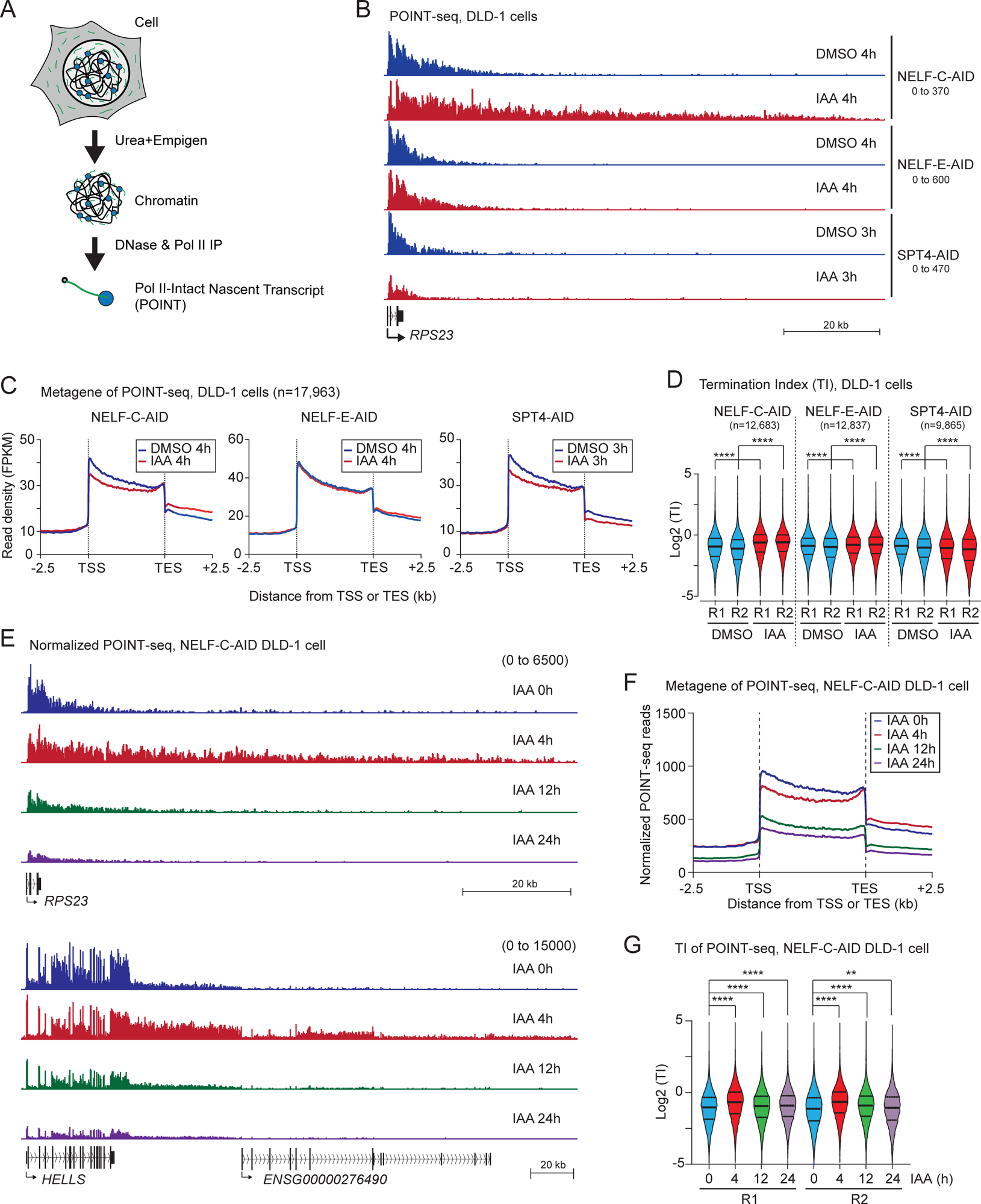
Acute NELF depletion fails to terminate Pol II transcription. (A) Schematic diagram of POINT-seq strategy. Chromatin was stringently isolated using Urea and Empigen detergent. Pol II intact nascent transcripts were precipitated with Pol II CTD antibody from DNA digested chromatin. (B) Example view of POINT-seq on *RPS23* gene in the indicated cell lines treated for 4h or 3h with DMSO or IAA. (C) Metagene analysis of POINT-seq on scaled TU -/+2.5kb in the indicated cell lines treated for 4h or 3h with DMSO or IAA. (D) Violin plots of Termination Index (TI) in the indicated cell lines treated for 4h or 3h with DMSO or IAA. Two biological replicates are shown. Statistical test: Wilcoxon signed-rank test. ****: p < 0.0001. (E) Example view of SIRV-normalized POINT-seq on *RPS23* and *HELLS* genes in NELF-C-AID DLD-1 cells (0, 4,12, and 24h IAA). (F) Metagene analysis of POINT-seq on scaled TU -/+2.5kb in NELF-C-AID DLD-1 cells (0, 4,12, and 24h IAA). (G) Violin plots of TI in NELF-C-AID DLD-1 cells (0, 4,12, and 24h IAA). Two biological replicates are shown. Statistical test: Wilcoxon signed-rank test. **: p < 0.01, ****: p < 0.0001.

As a defect in pre-mRNA splicing is known to also promote a transcription termination defect^50,51^, we investigated the effect of NELF-C protein on co-transcriptional splicing. Pladienolide B (PlaB), which inhibits early stages of splicing reaction through the binding to the U2 snRNP components SF3B1 protein, was used to evaluate co-transcriptional splicing. A 3h treatment of PlaB did not affect protein levels of NELF-C and Pol II in WCE of NELF-C-AID DLD-1 cells (Figure S3B). As previously reported, PlaB induced premature transcription termination (PTT) in pc genes, accompanying an inhibition of co-transcriptional splicing^14,52^. In addition, we found in these experiments that 3h IAA treatment extended PTT (Figure S3C). Notably, co-transcriptional constitutive splicing was unaffected by NELF-C depletion, although PlaB dramatically inhibits this splicing fraction (Figure S3D). These results indicate that the NELF complex contributes to Pol II transcription termination independently of co-transcriptional splicing.

NELF depletion caused a cell cycle arrest that started from 12h IAA treatment of NELF-C-AID DLD-1 cells (Figure 2D). This led us to analyze nascent RNAs at longer time points. While our POINT-seq data detected a Pol II termination defect of *RPS23* and *HELLS* genes at 4h IAA (Figures 3B-D), POINT-seq signals at 12h and 24h IAA were significantly reduced throughout their gene bodies (Figure 3E). Metagene analysis of POINT-seq confirmed a global termination defect after 4h IAA treatment and the global decrease in transcription after 12h and 24h IAA (Figure 3F). Additionally, compared to 0h IAA, higher TIs of POINT-seq were detected at 4h IAA, but not at 12h and 24h IAA (Figure 3G). Same as POINT-seq, the previously published PRO-seq data also detects transcription termination defect of *RPS23* gene after 4h IAA treatment of NELF-C-AID DLD-1 cells^10^ (Figure S3E). Notably the TI analysis of published PRO-seq data failed to detect the transient termination defect due to only faint signals downstream of PAS detected by PRO-seq, while only highly expressed pc genes displayed the transient termination defect (Figure S3F). Similarly to our POINT-seq, our TT-seq data displayed transcription termination defect of *RPS23* and *HELLS* genes (Figure S3G) and higher TIs (Figure S3H) after 4h IAA treatment of NELF-C-AID DLD-1 cells.

### NELF controls transcription termination via elongation rate, independently of promoter proximal Pol II pausing or 3’ poly(A) sites

Based on the TI, we classified genes into three categories: unchanged (UncTI, n=4,096), increased (IncTI, n=6,523), and decreased TI genes (DecTI, n=993) (Figure S4A). Metagenes of POINT-seq for IncTI genes, but not for UncTI genes, showed a clear Pol II transcription termination defect of pc genes at 4h IAA (Figure 4A). The fraction of genes in the IncTI category decreased over IAA treatment time (4h: 56%, 12h: 33%, 24h: 29%) (Figure 4B), consistent with the TI analysis (Figure 3G). We quantified such fractions in another human colorectal cancer line, HCT116 which expresses AID tagged CPSF73 and Xrn2 proteins, major transcription termination factors of pc genes, and found that the IncTI genes represented 81% and 66%, respectively (Figure 4B). These results indicate that NELF-C is a critical transcription terminator factor for pc genes in colorectal cancer cells.

**Figure 4.**
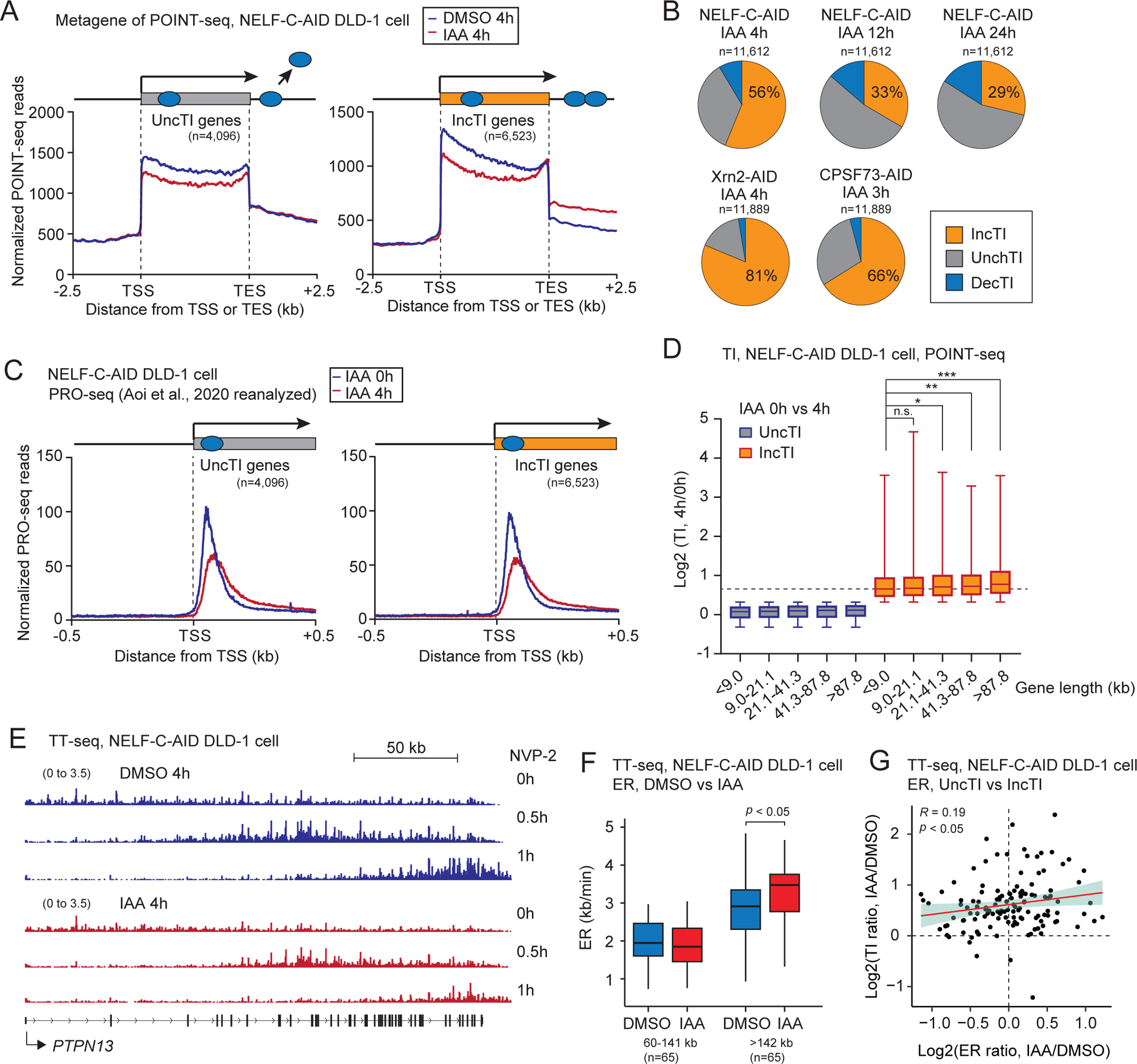
NELF controls transcription termination independently of promoter proximal Pol II pausing or 3’ poly(A) sites. (A) Metagene analysis of POINT-seq on scaled TU -/+2.5kb of Unchanged TI (UncTI) and Increased TI (IncTI) genes in NELF-C-AID DLD-1 cells (4h DMSO and IAA). (B) Pie charts of UncTI, IncTI, and Decreased TI (DecTI) genes in indicated AID-tagged cells. IAA incubation time is shown. (C) Metagene analysis of published PRO-seq on TSS -/+0.5kb of UncTI and IncTI genes. (D) Box plots of TI of UncTI and IncTI genes in NELF-C-AID DLD-1 cells (4h DMSO and IAA). Genes were classified by gene length as indicated. Statistical test: Wilcoxon rank sum test. N.s.: not significant, *: p < 0.05, **: p < 0.01, ***: p < 0.001. (E) Example view of TT-seq on *PTPN13* gene in the indicated cell line treated for 4h with DMSO or IAA, followed by NVP-2 treatment for 0, 0.5, or 1h. (F) Box plots of elongation rates (ER) derived from TT-seq. Data for short and long genes are shown. Statistical test: Wilcoxon rank sum test. (G) Scatter plot showing the correlation between ER and TI. The regression line (red) and the 95% confidence interval (green) are also shown. n = 129. Statistical method: Spearman correlation.

A previous study demonstrated that acute loss of NELF-C protein resulted in a perturbation of Pol II pausing at promoter-proximal sites shifting to downstream secondary pause sites^10^. To determine whether there is a link between genes with a perturbed Pol II pausing and genes with a transcription termination defect, we reanalysed the previously published PRO-seq data and detected similar profiles of Pol II secondary sites pause in both IncTI and UncTI categories (Figure 4C). Also, NELF-C occupancy at the promoter-proximal site detected by chromatin immunoprecipitation was unchanged between both categories (Figure S4B). To investigate promoter activity, we employed the ATAC-seq method at 4h and 24h IAA treatments. ATAC-seq profiles (Figure S4C) and their quantification (Figure S4D) on UncTI and IncTI genes upon NELF-C depletion demonstrated that ATAC-seq signals decrease similarly only at 24h IAA on promoter regions of both gene categories. Consistent with decreased POINT-seq after 24h IAA, this indicates that stable loss of NELF-C suppresses transcription initiation. Importantly, these observations demonstrate that NELF-C regulation of Pol II pausing at promoter-proximal sites is not connected to its effect on transcription termination.

To determine whether the IncTI and UncTI genes differ by the strength of their PAS, we analysed the adjacent sequences (-/+ 20 nt) of PAS in UncTI and IncTI genes. We failed to detect a significant difference between UncTI and IncTI genes (Figure S4E), even though the nucleotide compositions were slightly biased to a higher AT-rich in IncTI genes (Figure S4F).

We also investigated whether gene length correlates with TI change. Higher TI was detected in IncTI genes than in UncTI genes for all the gene length categories (Figure 4D). However, the longest gene category (>87.8kb) shows a higher TI, suggesting that Pol II transcription elongation activity is involved for an efficient transcription termination as longer genes are known to have a higher elongation rate (ER)^53^. In the nascent RNA analysis, a CDK9 inhibitor treatment showed transcription inhibition waves, indicating that Pol II pause-release step is affected by a CDK9 inhibitor, but not the elongation step^53^. Therefore, we measured the ER by performing TT-seq with a specific CDK9 inhibitor NVP-2 for 0.5h and 1h (Figure 4E). An example of the transcription inhibition waves, and the stronger effect on the inhibition waves upon NELF-C loss for 4h, are visible on *PTPN13* gene (Figure 4E). Due to the absence of TT-seq signal on the first 60kb region of genes following CDK9 inhibition, we focused on pc genes with reproducible detection of the inhibition waves (>60kb) for the global ER measurement (Figures 4F and S4G). This analysis demonstrated that NELF-C loss significantly increases Pol II transcription ER on genes longer than 142kb (+17% in median), whereas we did not observe this increase on shorter genes (60-141kb) (Figure 4F). Notably, our comparative analysis showed a positive correlation between increased Pol II speed and termination defect (Figure 4G). Our results suggest that NELF-C contributes to regulate Pol II speed to ensure an efficient transcription termination.

### NELF loss alters 3’end processing of pc genes

As NELF loss promotes a transcription termination defect, we investigated whether 3’end processing of pc genes is also affected. Nascent RNAs of RDH genes are co-transcriptionally cleaved by CPSF73 protein together with stem-loop binding protein (SLBP), FLASH, and U7 snRNP that binds to the histone downstream elements^54,55^. This results in production of non-polyadenylated histone mRNAs. Consistent with a previous study^56^, heatmap and quantification analysis of our nuclear pA+RNA-seq data (two biological replicates) demonstrated that the expression and the polyadenylation of RDH genes are dramatically up-regulated by NELF-C depletion (Figures 5A and S5A). For examples, pA+ RNA levels of *H2BC18* and *H2BC21* genes are increased up to ∼25 fold at 24h IAA (Figures 5B and S5B). In addition, our POINT-seq analysis detected Pol II transcription termination defects for RDH genes such as *H2BC21* (Figure 5A), as well as the other RDH genes in the cluster (Figure S5C), following NELF-C depletion. Metagene of POINT-seq displayed a clear termination defect of RDH genes at 4h IAA treatment (Figure S5D). Similarly, stable loss of NELF-C reduces over time the transcription levels of RDH genes (Figures 5B and 5C).

**Figure 5.**
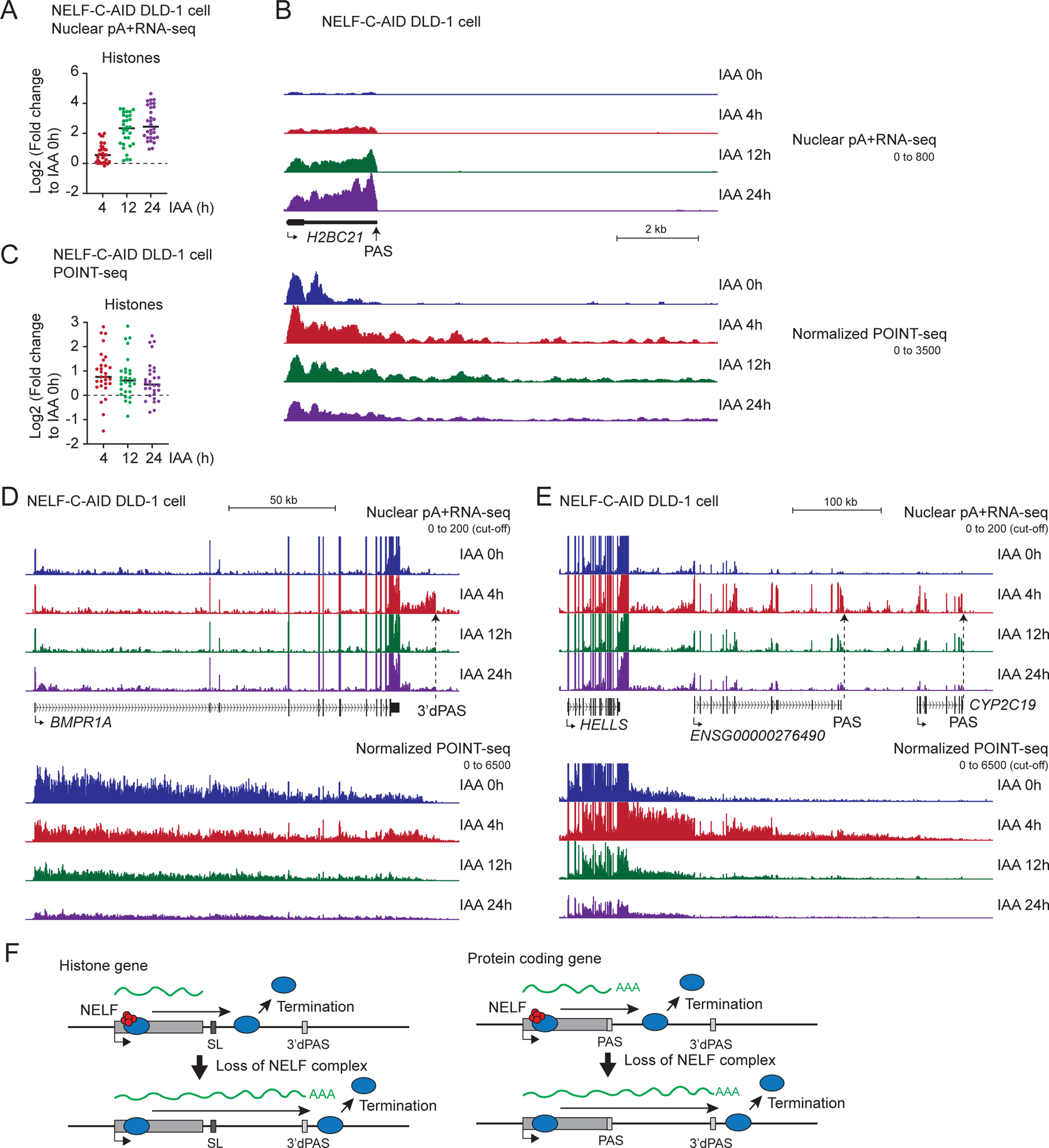
NELF loss alters 3’end processing of pc genes. (A) Log_2_ fold change of nuclear pA+ RNA-seq signals of RDH genes in NELF-C-AID DLD-1 cells (4, 12, and 24h IAA). (B) Example view of nuclear pA+ RNA-seq and SIRV-normalized POINT-seq on *H2BC21* gene in NELF-C-AID DLD-1 cells (0, 4,12, and 24h IAA). PAS is indicated with an arrow. (C) Log_2_ fold change of SIRV-normalized POINT-seq signals on RDH genes in NELF-C-AID DLD-1 cells (4, 12, and 24h IAA). (D) Example view of nuclear pA+ RNA-seq and SIRV-normalized POINT-seq on *BMPR1A* gene in NELF-C-AID DLD-1 cells (0, 4, 12, and 24h IAA). PAS is indicated with an arrow. (E) Example view of nuclear pA+ RNA-seq and SIRV-normalized POINT-seq on *HELLS* gene in NELF-C-AID DLD-1 cells (0, 4, 12, and 24h IAA). PASs are indicated with arrows. (F) Models of NELF-mediated Pol II transcription termination.

Extended readthrough transcript of the *BMPR1A* genes was detected by the POINT-seq analysis after 4h IAA. Under the same condition, we observed distal PAS (3’dPAS) usage located downstream of *BMPR1A* gene 3’end in our nuclear pA+ RNA-seq (Figure 5D). Similarly, 3’dPASs were also observed for *RPS23* and *CTPS1* genes in the extended transcription regions, following an acute loss of NELF (Figure S5E). For the *HELLS* gene, extended readthrough transcripts were spliced and polyadenylated at the downstream genes in the absence of NELF (Figure 5E).

These results indicate that loss of NELF protein suppresses non-PAS usage (such as stem loop, SL) and activates the 3’dPAS in RDH genes (Figure 5F). On the other hand, in pc genes, rapid loss of NELF protein activates 3’dPAS, but does not interfere with co-transcriptional constitutive splicing of readthrough transcripts.

### NELF loss causes Pol II transcription invasion into DNA replication initiation zone

DNA replication origin mapping using 5-ethynyl-2’-deoxyuridine (EdU)-labelled DNAs has demonstrated that nascent DNA synthesis is preferentially initiated in the intergenic regions of human U2OS and HeLa cell lines^57^. Consistent with this study, another technology to map replicative DNA polymerases, called Pu-seq (polymerase usage-sequencing) maps DNA replication initiation and termination zones over genomic regions with reciprocal demarcations in usage of leading and lagging strand polymerase in HCT116 human colorectal cancer cell^43^. Notably, Pu-seq technology also predicted that sites of replication initiation are generally located in intergenic regions around highly transcribed pc genes. This suggests that Pol II transcription and DNA replication are mutually exclusive in human cancer cells. In fact, perturbation of Pol II transcription termination causes T-R conflict, a major source of genomic stress in human cells^39,40^.

Based on the DNA replication initiation (RI) score (from 0.0363 to 265.5028) determined by quantification of the Pu-seq signals, 12,529 DNA RI zones were identified. To investigate Pol II transcription activity in the RI zone, we compared the POINT-seq signals of 4h IAA to that of 4h DMSO treatments in NELF-C-AID cells. As a representative example, POINT-seq signals in the *CD58* gene are displayed (Figure 6A). The mapped RI zone (score 54.6549) is located downstream of the *CD58* gene and upstream of *ATP1A1-AS1* lncRNA gene. Importantly both genes are transcriptionally active. The POINT-seq technology detected extended readthrough transcripts into the RI zone only in the absence of NELF-C protein (4h IAA) (Figure 6A). Similarly, readthrough transcripts were detected in the RI zone (score 22.7711) downstream of another example gene *FOCAD*, but again only in the absence of NELF-C protein (Figure S6A). *IFN* genes in the cluster in or near this RI region were encoded on the opposite strand of the *FOCAD* gene. However, *IFN* genes are not expressed. The generality of Pol II invasion in the RI regions after 4h IAA is demonstrated in metagene analysis (Figure 6B) and quantification of POINT-seq (Figure 6C). A stable loss of NELF-C displayed decreased POINT-seq signals in the RI regions (12h and 24h IAA), in agreement with the global decrease in transcription (Figure S6B). As expected from the lack of a transcription termination defect, acute loss of NELF-E and SPT4 proteins showed no significant increase of POINT-seq signals in the RI zones (Figures S6C and S6D). In addition, PlaB treatment dramatically reduced the POINT-seq signals in the RI zone (Figure S6E), since PlaB causes PTT in over 20% of expressed pc genes of HCT116 cells^14^.

**Figure 6.**
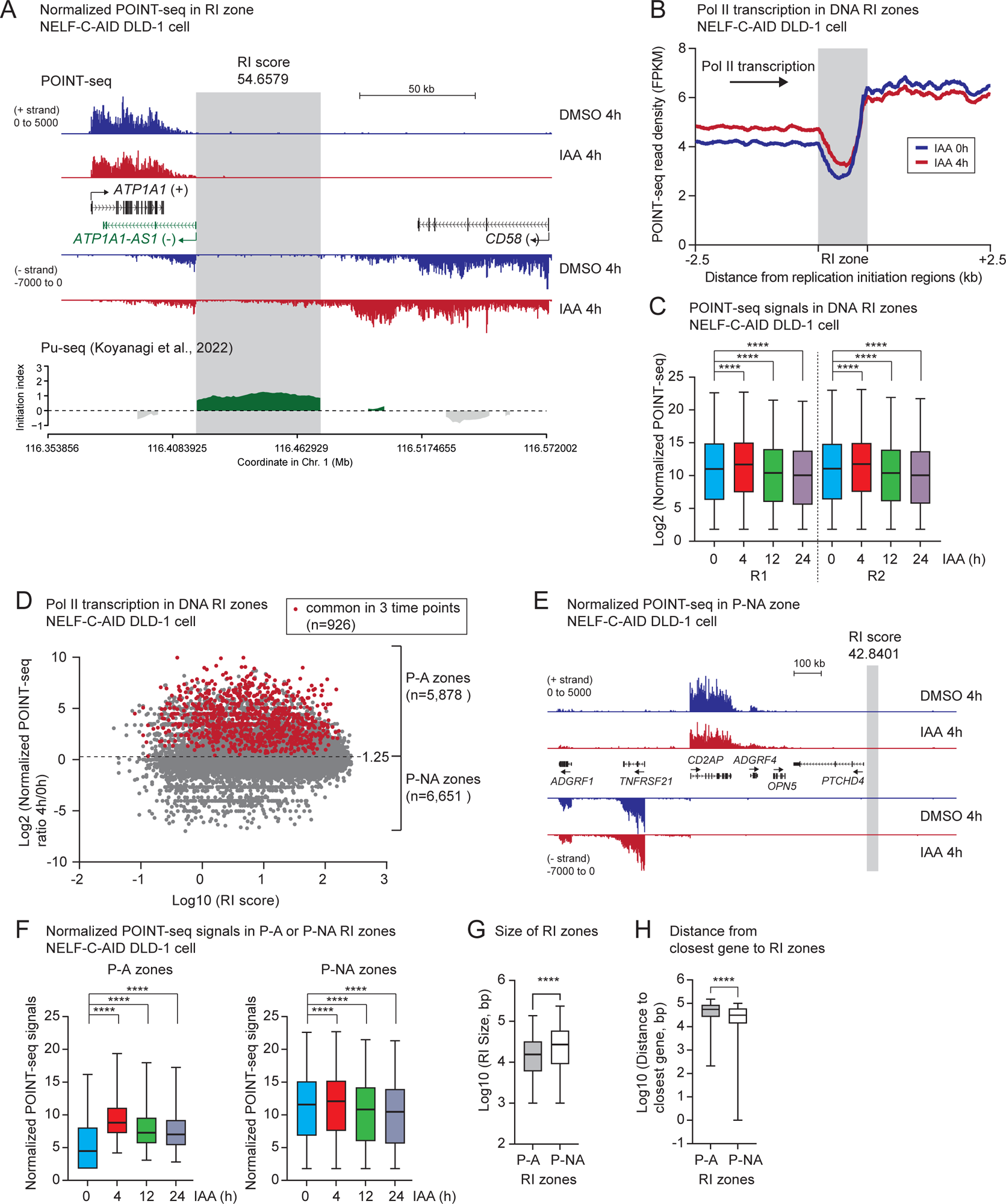
NELF loss causes Pol II transcription invasion into DNA replication initiation zone. (A) Example view of SIRV-normalized POINT-seq on *CD58* gene adjacent to a RI zone in NELF-C-AID DLD-1 cells (4h DMSO and IAA). DNA replication initiation (RI) zone is highlighted in gray. RI score: 54.6579. Re-analyzed Pu-seq data shows RI zones in green. (B) Metagene analysis of POINT-seq on scaled RI zones -/+2.5kb in NELF-C-AID DLD-1 cells (4h DMSO and IAA). (C) Box plots of SIRV-normalized POINT-seq signals in RI zones in NELF-C-AID DLD-1 cells (0, 4, 12, and 24h IAA). Two biological replicates are shown. Statistical test: Wilcoxon signed-rank test. ****: p < 0.0001. (D) Scatter plots of SIRV-normalized POINT-seq signals and RI score in NELF-C-AID DLD-1 cells (4h DMSO and IAA). Cut-off value is Log_2_(1.25) in SIRV-normalized POINT-seq. Pol II-affected (P-A) and not affected (P-NA) zones are classified as higher and lower than the cut-off value, respectively. RI zones that are above the cut-off in the three timepoints (4, 12, 24h IAA) are indicated in red. (E) Example view of SIRV-normalized POINT-seq on *CD2AP* gene adjacent to a P-NA zone in NELF-C-AID DLD-1 cells (4h DMSO and IAA). DNA RI zone is highlighted in gray. RI score: 42.8401. (F) Box plots of SIRV-normalized POINT-seq signals in P-A and P-NA RI zones in NELF-C-AID DLD-1 cells (0, 4, 12, and 24h IAA). Statistical test: Wilcoxon rank sum test. ****: p < 0.0001. (G) Size in bp of P-A or P-NA RI zones. Statistical test: Wilcoxon rank sum test. ****: p < 0.0001. (H) Distance in bp between the closest gene and the P-A and P-NA RI zones. Statistical test: Wilcoxon rank sum test. ****: p < 0.0001.

To extract the RI zones that are potentially perturbed by dysregulated Pol II transcription readthrough in the absence of NELF-C protein, we classified them into two categories using the ratio of 4h to 0h IAA POINT-seq signals (cut-off: 1.25): the Pol II-affected (P-A, n= 5,878) and not affected (P-NA, n=6,651) zones (Figure 6D). Common P-A zones between 4h, 12h, and 24h IAA are shown in red (n=926). This result suggests that not all the RI zones are occupied by elongating Pol II upon loss of NELF complex. For example, Pol II transcription termination was impaired in the representative pc gene *CD2AP*. However, the readthrough transcription did not reach to the neighbouring RI zone (score 42.8401) (Figure 6E). As an example of a P-NA zone, Pol II termination was not greatly perturbed for the *SPOPL* gene after acute loss of NELF-C protein, resulting in no POINT signal accumulation in the downstream RI zone (score 41.0896) (Figure S6F). Quantification of TI (4h vs 0h IAA) in NELF-C-AID DLD-1 cells demonstrated that pc genes adjacent to the P-A zones has a higher TI compared to genes close to P-NA zones (Figure S6G). As expected, POINT-seq signals were increased in P-A zones following NELF-C depletion, although those were unchanged in P-NA zones (Figure 6F). Notably POINT-seq signals in P-NA zones is higher than that in P-A zones. This may be due to the larger size of RI zones in P-NA category (Figure 6G). Additionally, we measured the distance between the closest gene and each RI zone, showing that P-A zones are more separated from transcription unit than P-NA zones (Figure 6H). This suggests that P-A zones need to be protected from Pol II transcription invasion. Indeed, 1,039 RI zones were identified in the category which has a RI score >10 and a ratio of POINT-seq (4h/0h) >2 fold. Perturbation of these zones are potentially associated with cell cycle arrest.

### Acute depletion of NELF-C protein causes global loss of DNA replication initiation factors on chromatin

Our results showing the deleterious effects of Pol II readthrough into P-A zones following NELF-C depletion led us to investigate the recruitment of DNA replication initiation factors to chromatin in the absence of NELF-C protein. The chromatin fractions of NELF-C-AID DLD-1 cells were isolated in a mild condition^58^ not using either Urea or Empigen detergent followed by proteomics. As expected, we observed a significant reduction of whole NELF complex components on chromatin after 4h IAA (Figures 7A and S7A and Supplementary Table 2), suggesting that this approach is able to assess NELF-dependent chromatin binding proteins. Notably the levels of several factors associated with DNA replication initiation, such as MCM3, MCM5, MCM1-3, MCM5, and CDT1 were slightly reduced on chromatin after 4h IAA (Figure 7B). CDT1 protein is required to fire the DNA replication origins in G1 phase, but is degraded in S phase to prevent aberrant DNA replication initiation^59^. To validate this result, we checked CDT1 protein in the chromatin by western blot analysis and found a reduction of CDT1 protein levels (Figure 7C). To increase the CTD1 protein levels, we used a small compound MLN-4924 that stabilizes CDT1 protein by inhibiting NEDD-8 activating enzyme^60^ for 24h after a 24h IAA pre-treatment in NELF-C-AID DLD-1 cells. As expected, the reduced cell population in S phase in the absence of NELF-C protein was largely recovered by MLN-4924 (Figure 7D).

**Figure 7.**
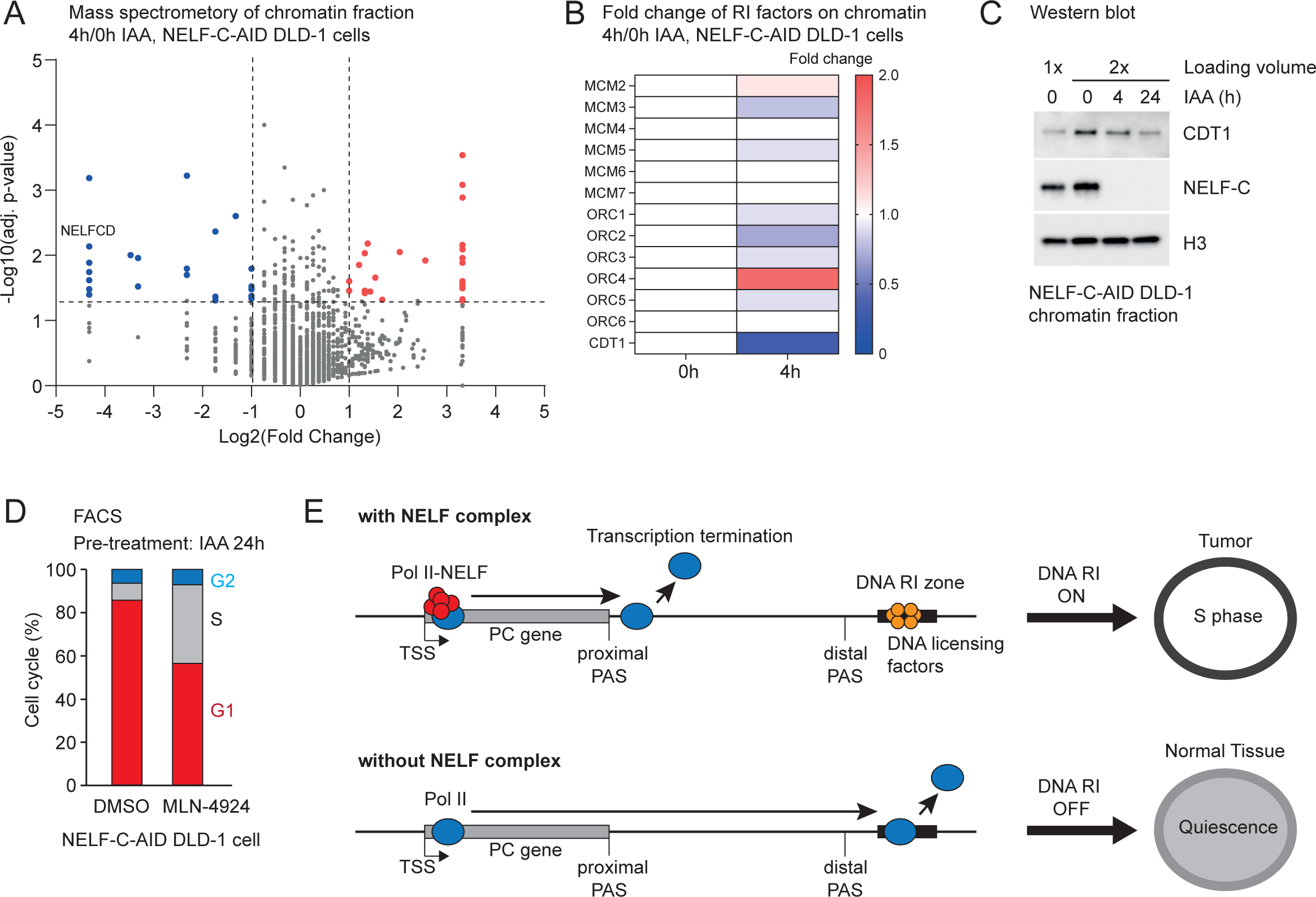
Acute depletion of NELF-C protein causes global loss of DNA replication initiation factors on chromatin. (A) Volcano plot for the log_2_ fold change (4h IAA / DMSO) of chromatin-associated proteins in NELF-C-AID DLD-1 cells. Proteins increased and decreased after 4h IAA are indicated in red and blue, respectively. NELF-C is indicated on the panel. (B) Heatmap for fold change of RI factors in NELF-C-AID DLD-1 cells (4h DMSO and IAA). (C) Western blot of chromatin fraction of NELF-C-AID DLD-1 cells (0, 4, and 24h IAA) using the indicated antibodies. (D) Cell cycle (%) of 24h DMSO or MLN-4924 treated NELF-C-AID DLD-1 cells. The cells were pre-treated with IAA for 24h. (E) Model of NELF-mediated transcription termination and DNA replication initiation. With NELF, Pol II transcription is terminated at proximal PAS before Pol II reaches DNA RI zone. Without NELF, Pol II transcription is extended to distal PAS and impair replication in the RI zone, resulting in cell quiescence.

Overall, our study suggests that perturbation of Pol II transcription termination caused by NELF depletion impairs DNA firing in particular RI zones called P-A zones (Figure 7E). A few hundreds ∼ thousands of RI zones may be perturbed by such transcription readthrough and potentially impair DNA replication licensing, leading to cell cycle arrest in G1 or early S phase (Figure 7E).

## DISCUSSION

Gene mutations, deletions, and amplifications disrupt transcriptional programs and transform normal cells into cancer cells^61^. Transcriptional dysregulation is established by altered expression levels of particular transcription and RNA processing factors. Therefore, cancer cells can be highly dependent on such transcription-associated factors. We found that transcripts of CPA factors are highly expressed in various tumors including COAD (Figure 1B). This observation suggests that CPA-dependent Pol II transcription termination is more efficient in tumor than in normal tissue. Indeed, JTE-607, an inhibitor of CPSF73 leads several cancer cells to apoptosis^62–64^. We additionally found that RNA expression of *NELFCD* is up-regulated especially in colorectal tumors (Figures 1E and S1B). NELF-C expression is required for cell proliferation (Figure 2E), in agreement with a previous study showing that siRNA knockdown of NELF-CD protein suppresses tumor growth in mouse^65^. Therefore, NELF proteins could potentially be potent targets of anti-cancer drugs. Indeed, knockout of *NELF-B* gene impairs cell proliferation and differentiation in mouse embryonic stem cells^49^. Transposon insertion into the *NELF-A* gene in fly embryos leads to a developmental failure^66^. Taken together, these observations support the view that the NELF complex has an evolutionary conserved role in cell proliferation.

NELFs were originally identified as negative transcription factors together with DSIF *in vitro*^7^. Our present study demonstrates that acute loss of the NELF complex perturbed Pol II transcription termination in colorectal cancer cells, while depletion of the DSIF component SPT4 had no effect. In this study, we did not investigate the function of the other DSIF component, SPT5 on transcription since it was shown to stabilizes the Pol II complex^11^. We conclude that the NELF complex contributes to Pol II transcription termination independently of DSIF.

Loss of NELF proteins shifts Pol II pausing from a promoter-proximal site to its downstream site called 2^nd^ Pause^10^. Our study did not detect a correlation between transcription termination defects and shifted promoter-proximal Pol II pausing. This indicates that the effect of NELF-C loss at the 3’end of pc genes is independent of its effect at the 5’end. In addition, transcription termination readthrough is not mediated via a pre-mRNA splicing defect as NELF-C loss does not inhibit RNA co-transcriptional splicing (Figure S3D). It still remains unclear how only transcription termination of a subset of pc genes is affected by the loss of NELF complex. Importantly, we observed that especially in long genes, IncTI genes have a higher Pol II elongation rate compared to UncTI genes after NELF loss (Figure 4G), supporting the hypothesis that the termination defect of IncTI genes is mediated by an increased Pol II speed. However, such transcription termination defect was also observed in some short genes including RDH and *RPS23* genes. Therefore, we consider the following non-mutually exclusive models. (1) Faster Pol II elongation is established in the absence of the NELF complex and so may interfere with the recognition of the proximal PAS or may fail to be caught by a torpedo activity of Xrn2, especially in long pc genes. (2) Decreased recruitment of particular CPA factors regulating proximal PAS. Notably we did find that the chromatin levels of a few CPA factors, such as CPSF6, CPSF7, CSTF1, and WDR33, are also significantly reduced in the absence of NELF proteins (Figure S7B). This reduction may contribute to alternative PAS usage followed by extended transcription readthrough. (3) Biochemical approaches have previously detected an interaction between the Integrator complex and the NELF complex in in HeLa S3 cells^67^, suggesting that this interaction has a role on transcription termination since loss of the Integrator complex attenuates pc gene transcription^17,68^.

T-R conflict is a major source of DNA damage. Previous studies have demonstrated that collisions can occur between running DNA replisome and Pol II (head to head) or Pol II and/or Pol II-derived RNA:DNA hybrid (head to tail)^69,70^. An siRNA screen demonstrated that loss of WDR33, a CPA factor, causes global perturbation of Pol II transcription termination and a higher cell sensitivity to DNA replication stress from T-R conflict^39^. This finding suggests that Pol II transcription activity needs to be restricted to certain regions by transcription termination. Loss of CPSF73 and Xrn2 proteins severely disrupts transcription termination of pc genes^14,71,72^ (Figure 4E). Therefore, inhibition or depletion of these proteins induces a catastrophic DNA damage and cell death^63,73^. Additionally, our previous study demonstrated that siRNA-depletion of SPT6 protein globally upregulates lncRNA transcription and perturbs its termination^40^. This leads to T-R conflict followed by cell cycle arrest in G0 phase and then cellular senescence. In this study, we propose a novel T-R conflict that occurs between Pol II and DNA replication origin firing in G1 or early S phase. Pol II transcription termination was globally impaired by depletion of the NELF complex, but was limited over time to a subset of genes (∼3,300 pc genes affected after 24h IAA). This limited defect is still sufficient to sustain cell cycle arrest without inducing cell death, indicating that cells can still enter quiescence following NELF loss. In fact, normal tissues that contains non-replicative somatic cells express NELF-C protein at a lower level^65^. Our study now provides insight into T-R conflict in replicative cells that contributes to establish cell quiescence.

## AUTHOR CONTRIBUTIONS

CN and MT conducted the bioinformatic analyses. QF and TN performed the molecular biology and POINT-seq experiments. YD performed FACS analysis. YA and AS performed TT-seq analysis for transcription elongation rate. HK provided valuable Pol II antibodies. TN and MT designed the project. CN, QF, YA, AS, YD, MT, and TN co-wrote the paper.

## ACKNOWLEDGMENTS

We thank Dr. Mikita Suyama and Dr. Chie Kikutake for help with the computational analyses, Dr. Rebekah Jukes-Jones for help with the proteomics analysis, and Dr. Nick J Proudfoot and Dr. Kinga Kamieniarz-Gdula for critical comments on this paper. This work was supported by funding to TN (MEXT/JSPS Kakenhi (JP19K24692) and JST FOREST (JPMJFR2050), Astellas Foundation for Research on Metabolic Disorders, Daiichi Sankyo Foundation of Life Science, The Ichiro Kanehara Foundation for Promotion of Medical Sciences and Medical Care, The Mitsubishi Foundation, The Mochida Memorial Foundation for Medical and Pharmaceutical Research, The Naito Foundation, The NOVARTIS Foundation (Japan) for the Promotion of Science, Princess Takamatsu Cancer Research Fund, The Shinnihon Foundation of Advanced Medical Treatment Research, The Sumitomo Foundation, and The Uehara Memorial Foundation), MT (MRC (MR/W007002/1)), YD (MEXT/JSPS Kakenhi (JP23H02463), JST FOREST (JPMJFR204X)), AS (NIH (R35CA197569)) and Fellowship to CN (JST SPRING (JPMJSP2136)).

