## Supplementary information for "NELF coordinates Pol II transcription termination and DNA replication initiation"

[8]Lead author

#### STAR METHODS

KEY RESOURCES TABLE

| REAGENT OR RESOURCE | SOURCE | IDENTIFIER |
|---|---|---|
| Reagent | | |
| Auxin (IAA) | Sigma | Cat# I2886 |
| Dulbecco's Modified Eagle Medium (DMEM - High glucose) | Nacalai Tesque | Cat# 29113-53 |
| Empigen ~30% | Sigma | Cat# 30326 |
| Turbo DNase | Thermo Fisher Scientific | Cat# AM2239 |
| Benzonase | Millipore | Cat# E1014 |
| RiboLock RNase inhibitor | Thermo Fisher Scientific | Cat# EO0381 |
| Tri Reagent | Cosmo Bio | Cat# TR118 |
| Dynabeads M280 Sheep Anti-mouse IgG | Thermo Fisher Scientific | Cat# 11202D |
| Dimethyl sulfoxide (DMSO) | Sigma | Cat# D8418 |
| Pladienolide B | Santa Cruz | Cat# SC-391691 |
| MLN-4924 | Cell Signaling | Cat# 85923 |
| SPRISelect reagent | Beckman Coulter | Cat# B23317 |
| Spike-in SIRV-Set2 | Lexogen | Cat# 050.0 |
| Chemi-Lumi One Super | Nacalai Tesque | Cat# 02230-30 |
| NuPAGE LDS Sample Buffer (4x) | Thermo Fisher Scientific | Cat# NP0007 |
| 4-Thiouridine | Sigma | Cat# T4509 |
| NVP-2 | Med Chem Express | Cat# HY-12214A |
| MTSEA-biotin-XX | Biotium | Cat# 90066 |
| Dynabeads MyOne Streptavidin C1 | Thermo Fisher Scientific | Cat# 65001 |
| Antibodies | | |
| Mouse monoclonal anti-Pol II CTD, Total | Previous study[1] | MABI601 for IP, Available from Kimura Lab by request. |
| Mouse monoclonal anti-POLR2A (N-terminal) | Santa Cruz | Cat# sc-55492, for western blot |

| | | |
|---|---|---|
| Mouse monoclonal anti-RPA194 (Pol I) | Santa Cruz | Cat# sc-48385 |
| Rabbit monoclonal anti-NELFB (COBRA1) | Abcam | Cat# ab167401 |
| Mouse monoclonal anti-NELFA | Santa Cruz | Cat# sc-365004 |
| Rabbit monoclonal anti-NELFE | Abcam | Cat# ab170104 |
| Rabbit monoclonal anti-NELFC/D (TH1L) | Cell Signaling | Cat# 12265 |
| Mouse monoclonal anti-CDT1 | Santa Cruz | Cat# sc-365305 |
| Rabbit monoclonal anti-SPT4 | Cell Signaling | Cat# 64828 |
| Mouse monoclonal anti-Tubulin (clone DM1A) | Sigma | Cat# T6199 |
| Rabbit polyclonal anti-P57 (Kip2) | Cell Signaling | Cat# 2557 |
| Rabbit monoclonal anti-p21 (Cip1) | Cell Signaling | Cat# 2947 |
| Mouse monoclonal anti-H3 | Active Motif | Cat# 39763 |
| Mouse monoclonal anti-Histone H2A.X | Santa Cruz | Cat# sc-517336 |
| Rabbit monoclonal anti-Histone H2A.X Ser139p | Cell Signaling | Cat# 2577S |
| Goat anti-Mouse IgG H&L (HRP) | Abcam | Cat# ab205719 |
| Goat anti-Rabbit IgG H&L (HRP) | Abcam | Cat# ab205718 |
| Deposited Data | | |
| Raw sequencing data | This study | GEO: GSE253121 |
| Re-analysed NELF-C ChIP-seq data | Previous study[2] | GEO: GSE144786 |
| Re-analysed PRO-seq data | Previous study[2] | GEO: GSE144786 |

| | | |
|---|---|---|
| Re-analysed POINT-seq data (CPSF73, Xrn2) | Previous study[3] | GEO: GSE159326 |
| Processed TCGA and GTEx count data | Previous study[4] | Figshare https://doi.org/10.6084/m9.figshare.5330539 |
| Cell lines | | |
| Parental DLD-1 OsTIR1 cell (human) | Previous study[2] | Available from Shilatifard Lab by request. |
| NELF-C-AID DLD-1 OsTIR1 cell #7-10B (human) | Previous study[2] | Available from Shilatifard Lab by request. |
| NELF-E-AID DLD-1 OsTIR1 cell #20-1B (human) | Previous study[2] | Available from Shilatifard Lab by request. |
| SPT4-AID DLD-1 OsTIR1 cell #5-1D (human) | Previous study[5] | Available from Shilatifard Lab by request. |
| Gels | | |
| Novex 6% TBE gel, 12 well | Invitrogen | Cat# EC62652BOX |
| 4-15% Mini-PROTEAN Precast Protein Gels, 12 well | BioRad | Cat# 4561085 |
| Kits | | |
| NEBNext Ultra II Directional RNA library prep kit for Illumina | NEB | Cat# E7760S<br>(Note: this kit for POINT-seq) |
| Direct-zol RNA microprep | Zymo Research | Cat# R2061 |
| NEBNext PolyA mRNA Magnet isolation module | NEB | Cat# E7490S |
| ATAC-Seq Kit | Active Motif | Cat# 53150 |

| | | |
|---|---|---|
| Click-iT Plus EdU Alexa Fluor488 Flow Cytometry Assay Kit | Thermo Fisher Scientific | Cat# C10633 |
| NEBNext Magnesium RNA Fragmentation Module | NEB | Cat# E6150 |
| NEBNext rRNA Depletion Kit v2 | NEB | Cat# E7400 |
| Software and Algorithms | | |
| FastQC (v0.11.5) | https://www.bioinformatics.babraham.ac.uk/projects/fastqc/ | N/A |
| Cutadapt (v1.18) | https://cutadapt.readthedocs.io/en/stable/ | Martin M. 2011[6] |
| TrimGalore (v0.4.4) | https://www.bioinformatics.babraham.ac.uk/projects/trim_galore/ | N/A |
| STAR (v2.7.0) | https://github.com/alexdobin/STAR | Dobin et al., 2013[7] |
| SAMtools (v1.9) | http://www.htslib.org/ | Li et al., 2009[8] |
| BEDtools (v2.29.2) | https://bedtools.readthedocs.io/en/latest/ | Quinlan et al., 2010[9] |
| deepTools2 (v3.4.2) | https://deeptools.readthedocs.io/en/develop/ | Ramirez et al 2016[10] |
| bedGraphToBigWig (v2.10) | https://hgdownload.soe.ucsc.edu/admin/exe/linux.x86_64/ | N/A |
| Salmon (v1.2.1) | https://salmon.readthedocs.io/en/latest/index.html | Patro et al., 2017[11] |
| DESeq2 (v1.30.1) | https://bioconductor.org/packages/release/bioc/html/DESeq2.html | Love et al., 2014[12] |
| Apgelm (v1.18.0) | https://bioconductor.org/packages/release/bioc/html/apeglm.html | Zhu et al., 2019[13] |

| | | |
|---|---|---|
| PEAKS XPro | https://www.bioinfor.com/peaks-studio/ | N/A |
| Scaffold 5 Viewer (v5.3.1) | https://www.proteomesoftware.com/products/scaffold-5 | N/A |
| dplyr (v1.1.2) | https://cran.r-project.org/web/packages/dplyr/index.html | N/A |
| ggplot2 (v3.4.3) | https://cran.r-project.org/web/packages/ggplot2/index.html | N/A |
| tidyr (v1.3.0) | https://cran.r-project.org/web/packages/tidyr/index.html | N/A |
| stringr (v1.5.0) | https://cran.r-project.org/web/packages/stringr/index.html | N/A |
| Enhanced Volcano (v1.14.0) | https://github.com/kevinblighe/EnhancedVolcano. | N/A |
| Harmonizome3.0 | https://maayanlab.cloud/Harmonizome/gene_set/RNA+Polymerase+II+Transcription/Reactome+Pathways | Rouillard et al., 2016[14] |
| proRate | https://github.com/yuabrahamliu/proRate | N/A |

#### RESOURCE AVAILABILITY

#### LEAD CONTACT

Further information and requests for resources and reagents should be directed to Takayuki Nojima.

#### MATERIAL AVAILABILITY

All reagents in this study are commercially available except for anti-Pol II CTD antibody which is given by Dr. Hiroshi Kimura.

#### DATA AND CODE AVAILABILITY

Raw and processed data of POINT-seq, ATAC-seq, TT-seq, and nuclear pA+RNA-seq generated in this study are deposited in NCBI GEO and accessible through GSE253121. Raw image data associated with this study are available from Mendeley (available when published). All code supporting POINT analyses are available on request. Published data reanalysed in this study can be found at GEO (see Key Resource Table)

#### METHOD DETAILS

##### Cell culture

Details of DLD-1 cell lines (NELFC-AID DLD-1, NELFE-AID DLD-1, SPT4-AID DLD-1, and the parental DLD-1 OsTIR1) were previously published[2,5]. Cells were cultured in high glucose Dulbecco's Modified Eagle's Medium (DMEM) with 10% fetal bovine serum (FBS) and penicillin/streptomycin. All the cells were maintained in a humidified incubator at 5% CO2, 37ºC.

##### IAA treatment and cell count

AID-tagged protein degradation was induced by adding 500 µM auxin (IAA) to the culture media. In the recovery experiment, cells were washed twice with pre-warmed PBS before adding fresh media to ensure the complete removal of IAA.  Cell counting was completed using the Countess 3 FL system (Thermo Fisher Scientific).

##### Western blot

The primary antibodies used in this study are listed in the key resource table. Goat anti-Mouse IgG H&L (HRP) and Goat anti-Rabbit IgG H&L (HRP) were used as secondary antibodies. Chemi-Lumi

One Super was used for the chemiluminescence. Images were captured by the Chemidoc Touch V3 system (Bio-Rad).

**Polymerase intact nascent transcript sequencing (POINT-seq)**

Protocol was slightly modified in this study[3]. Typically, 10 million cells were lysed with 4 mL ice-cold hypotonic buffer (10 mM Tris-Cl pH 7.5, 10 mM NaCl, 2.5 mM $MgCl_2$, 0.5% NP-40) on ice for 5 min. The cell lysate was carefully underlaid with 1 mL sucrose cushion buffer (hypotonic buffer with 10% (w/v) sucrose). The nuclear fraction was pelleted by centrifugation (300 g, 5 min at 4 $^{o}$C), then washed with ice-cold hypotonic buffer (without NP-40) once. The purified nuclear fraction was first resuspended in 300 μL (200~500 μL, depending on the pellet size) ice-chilled NUN1 buffer (20 mM Tris-HCl pH 7.9, 75 mM NaCl, 0.5 mM EDTA pH 8.0, and 50% glycerol) and then lysed in 3 mL (at least 10-fold volume of NUN1) ice-cold NUN2 buffer (20 mM HEPES-KOH pH 7.6, 300 mM NaCl, 0.2 mM EDTA pH 8.0, 7.5 mM $MgCl_2$, 1% NP-40, 1 M Urea, 3% Empigen BB, 1x Protease inhibitor Complete EDTA-free, and 1x PhosSTOP). Note: The cotton candy-like chromatin should appear within a few minutes. The chromatin was then pelleted and washed with ice-cold PBS once (300 g, 1 min at 4 $^{o}$C). Chromatin-bound Pol II was released by DNase digestion in the reaction mixture (10 mM Tris-HCl pH 7.5, 400 mM NaCl, 100 mM $MnCl_2$, 2 Units/ μL RiboLock, and 0.2 Units/μL Turbo DNase) at 37$^{o}$C for 15 min in the ThemoMixer C (Eppendorf) at 1200rpm. EDTA pH 8.0 (final 1mM) was used to stop the DNase reaction, and the supernatant was collected by high-speed centrifugation (14,000g, 10 min at 4$^{o}$C). A 5-fold volume of ice-cold NET-2E buffer (50 mM Tris-HCl pH 7.4, 150 mM NaCl, 0.05 % NP-40, and 3% Empigen BB) was used for the dilution of the supernatant.

The 10 μg of anti-Pol II antibody MABI601 were mixed with 200 μL anti-mouse IgG Dynabeads solution in 1 mL NET-2 buffer (NET-2E without Empigen) and incubated at 4 $^{o}$C with mild rotation for at least 1 hour. The conjugated beads were washed with NET-2 buffer once and resuspended in 200 μL NET-2E buffer. The resuspended beads were added to the diluted supernatant of chromatin for at least 1 hours and then incubated at 4$^{o}$C with mild rotation (Note: Avoid overnight incubation). After the incubation, beads were washed with 1 mL of ice-cold NET-2E buffer at least 6 times. Pol II-bound nascent RNAs (POINTs) were extracted from the beads using Tri Reagent and subsequently purified by Direct-zol kit following the manufacturer's protocol.

Equal amount of SIRV spike-in oligos were added to each POINTs. The POINTs were fragmented to 150-300 nucleotides according to the NEBNext Ultra II Directional RNA library prep kit. Library preparation was completed following the manufacturer's protocol. The libraries were applied to Illumina PE150 NovaSeq6000 service (Novogene). Each POINT-seq sample should have more than 30 million reads.

**Transient Transcriptome sequencing (TT-seq)**

TT-seq samples were prepared as previously described[15] with the following modifications. NELF-C-AID DLD-1 cells were treated with DMSO or 500 µM IAA for 4h. Cells were then treated with CDK9 inhibitor NVP-2 at 250 nM for 0, 0.5 or 1h, followed by in vivo RNA labelling with 500 µM 4-Thiouridine for 10 min. Total RNA was extracted using TRIzol. ~200 µg total RNA was fragmented at 94°C for 2 min using NEBNext Magnesium RNA Fragmentation Module. Biotinylation of the fragmented 4sU-labeled RNA was performed using 33 µg/ml MTSEA-biotin-XX in the presence of 20% DMF for 1.5h at r.t. The biotinylated RNA was purified using Dynabeads MyOne Streptavidin C1 and eluted in 100 mM DTT. Library preparation with ~100 ng of the enriched RNA was performed using NEBNext rRNA Depletion Kit v2 and NEBNext NEBNext Ultra II Directional RNA Library Prep Kit. DNA libraries were sequenced on NovaSeq 6000 (Illumina).

**Nuclear-pA+ RNA-seq**

Part of the POINT-seq nuclear fraction was used for nuclear pA+ RNA extraction. Nuclear RNAs were extracted from DLD-1 cells using Tri Reagent and subsequently purified by Direct-zol kit following the manufacturer's protocol. NEBNext PolyA mRNA Magnet isolation module for 5 µg of nuclear RNA was employed to enrich the nuclear pA+ RNAs. The 100 ng of nuclear pA+ RNAs are processed as POINT for the sequencing library preparation.

**ATAC-seq**

ATAC-seq samples were prepared from 1 million NELF-C-AID DLD-1 cells by following the manufacturer's protocol (ATAC-seq kit, Active Motif). IAA (500 µM) was incubated for 0h, 4h, and 24h to degrade NELF-C protein. The libraries were applied to Illumina PE150 NovaSeq6000 service (Novogene).

**Fluorescence activated cell sorter (FACS) analysis**

To examine the effect of recovery from the degradation of NELF-C protein on cell cycle, at 24h time point after 24h IAA (500 µM) treatment in NELF-C-AID DLD-1 cells, IAA was washed off by rinsing cells on culture dishes with media twice. These cells were further incubated with media without IAA for 24h or 48h. Alternatively, to examine the effect of MLN-4924, at the same time point, 300 nM of MLN-4924 or DMSO were added to cell culture media and further incubated for 24h or 48h.

Next, cells were incubated with media containing 10 µM of 5-ethynil-2'-deoxyuridine (EdU) for 1.5h. After adding 0.1% sodium azide to media, cells were trypsinized and harvested. Using Click-iT Plus EdU Alexa Fluor488 Flow Cytometry Assay Kit, 3 x $10^6$ cells were subjected to fixation, permeabilization and the detection of EdU. After these treatments, cells were then incubated with 40 µg/mL of propidium iodide and 0.1 mg/mL RNase A for 30 m at room temperature and analysed using BD Accuri C6 Plus (BD bioscience). Data were processed using FlowJo Software (v10.10, BD bioscience).

**Chromatin fraction for quantitative mass spectrometry**

Protocol was derived from a previous study[16] with minor modifications. In brief, nuclear extraction was obtained by pipetting (20 times) instead of homogenizing, and the subsequent lysis step was extended to 30 min. The chromatin pellet was digested using Benzonase-containing buffer (150 mM HEPES pH 7.9, 1.5 mM $MgCl_2$, 150 mM potassium acetate, and 100 Units/mL Benzonase) and then denatured in 1x LDS buffer for 10 min at 70°C.

**LC-MS/MS**

Proteins were fractionated by SDS-PAGE on 12% acrylamide gels. Gel bands were cut-out and subjected to in-gel trypsin enzymatic digestion. Resulting peptides were dissolved in a solution containing 0.1% trifluoroacetic acid and 2% acetonitrile and analyzed by an Orbitrap Exploris 240 mass spectrometer with the Field Asymmetric Ion Mobility Spectrometry (FAIMS) Pro device (Thermo Fisher Scientific, Waltham, MA) coupled with a Vanquish Neo UHPLC system (Thermo Fisher Scientific, Waltham, MA). Peptide separation was performed with a NANO HPLC Capillary Column (internal diameter, 0.075 mm; length, 12.5 cm; tip internal diameter, 0.05 mm) packed with 3-μm C18 column (Nikkyo Technos, Japan). The column was placed on the column oven compartment and maintained at a temperature of 40 °C. The mobile phases consisted of 0.1% formic acid (A) and 80% acetonitrile in water (B). Peptides were eluted with a gradient of 4_40% B for 20 min at a flow rate of 300 nL/min.

Full mass spectra were acquired in the mass-to-charge (m/z) range of 300 to 1500, achieving a resolution of 60,000 at m/z 200. The FAIMS compensation voltage (CV) settings were -45V and -65V. Higher Energy Collisional Dissociation (HCD) spectra were collected automatically in data-dependent scan mode, utilizing a dynamic exclusion feature. The total cycle times for data-dependent scanning were 1.5 seconds at CV=-45V and 1.0 second at CV=-65V, respectively. For full MS, the normalized Automatic Gain Control (AGC) was set to 300%, and for HCD MS/MS, it was set to Standard Mode. The normalized collision energy was set at 30%. Before each run, a lock mass correction using the EASY-IC ion source assembly was performed.

#### QUANTIFICATION AND STATISTICAL ANALYSIS

**TCGA and GTEx analyses**

The processed TCGA and GTEx data was downloaded from a previous study[4]. In this study, only paired-end samples in normal tissue (N) and tumor (T) types which have more than 95 samples were employed. Final sample number after screening is indicated in Table 1. For comparison of normalized expression across N and T types, RSEM FPKM data were used in gglot2. For differential expression analysis in DESeq2, RSEM count data were used for only the genes with a fold change (T/N) < -1.5 or > 1.5 and an adjusted p-value below 0.05. Data were visualized by EnhancedVolcano. RSEM counts of genes associated with RNA polymerase II transcription were analyzed in Harmonizome 3.0.

**Gene Annotation**

The list of protein-coding genes was extracted from the Gencode V41 annotation, based on the hg38 version of the human genome. To obtain the annotation of the highest transcript isoform expressed for each protein-coding gene, transcripts expression was quantified with Salmon on the biological replicates of nuclear pA+ RNA-seq treated with DMSO, resulting in a list of 17,963 expressed protein-coding genes. The list of exons used for the splicing analysis was obtained by extracting the location of exons in Gencode V41 annotation from the most expressed transcript isoform of each of the 17,963 expressed protein-coding genes.

**POINT-seq analysis**

Adapters were trimmed with Cutadapt in paired-end mode with the following options: --minimum-length 10 -q 15,10 -j 16 -A GATCGTCGGACTGTAGAACTCTGAAC -a AGATCGGAAGAGCACACGTCTGAACTCCAGTCAC. Trimmed reads were mapped to the human GRCh38.p13 reference sequence or the SIRV-Set 2: SIRV isoform Mix E0 sequences (See Lexogen website) with STAR and the parameters: --runThreadN 16 -- readFilesCommand gunzip -c -k -- limitBAMsortRAM 20000000000 --outFilterMultimapNmax 1 --outFilterScoreMin 10 --outSAMtype BAM SortedByCoordinate. SAMtools was used to retain the properly paired and mapped reads (-f 3). For non-spiked POINT-seq, FPKM-normalized bigwig files were created with deepTools2 bamCoverage tool with the parameters -bs 10 -p max –filterRNAstrand forward (or reverse) --normalizeUsing RPKM.

For SIRV spiked POINT-seq, SAMtools view with the –s option was used to subsample all the bam files to the bam file containing the lowest number of reads and then to create strand-specific BAM files. The normalization factor was calculated as: (number of SIRV reads) / (number of human reads + number of SIRV reads). Spiked normalized bedGraph files were created with BEDtools genomecov and the options: -bg -split –scale (1 / normalization factor). Bigwig files were generated from the BedGraph files with the bedGraphToBigWig tool.

**RNA-seq analysis**

Adapters were trimmed with Cutadapt in paired-end mode with the following options: --minimum-length 10 -q 15,10 -j 16 -A GATCGTCGGACTGTAGAACTCTGAAC -a AGATCGGAAGAGCACACGTCTGAACTCCAGTCAC. The remaining rRNA reads were removed by mapping the reads to the rRNA genes defined in the human ribosomal DNA complete repeating unit (GenBank: U13369.1) with STAR and the parameters --runThreadN 16 --readFilesCommand gunzip -c -k --outReadsUnmapped Fastx --limitBAMsortRAM 20000000000--outSAMtype BAM SortedByCoordinate. The unmapped reads were mapped to the human GRCh38.p13 reference sequence with STAR and the parameters: --readFilesCommand gunzip -c -k --limitBAMsortRAM 20000000000 --outFilterType BySJout --outFilterMultimapNmax 20 --alignSJoverhangMin 8 --alignSJDBoverhangMin 1 --outFilterMismatchNmax 999 --outFilterMismatchNoverReadLmax 0.04 --alignIntronMin 20 --alignIntronMax 1000000 --alignMatesGapMax 1000000 --quantMode GeneCounts --outSAMtype BAM SortedByCoordinate. SAMtools was used to retain the properly paired and mapped reads (-f 3) and to create strand-specific BAM files. FPKM-normalized bigwig files were created with deepTools2 bamCoverage tool with the parameters -bs 10 -p max –normalizeUsing RPKM.

For differential expression analysis, the outputs of STAR –quantMode GeneCounts were imported DESeq2 and apeglm keeping only the genes with a fold change $< -2$ or $> 2$ and an adjusted p-value below 0.05.

**PRO-seq analysis**

Adapters were trimmed with Cutadapt with the following options: --minimum-length 10 -q 15, 10 -j 16 -a AGATCGGAAGAGCACACGTCTGAACTCCAGTCA. The rRNA reads were removed by mapping the reads to the rRNA genes defined in human and in *Drosophila melanogaster* with STAR and the parameters --runThreadN 16 --readFilesCommand gunzip -c -k --outReadsUnmapped Fastx --limitBAMsortRAM 20000000000--outSAMtype BAM SortedByCoordinate. The unmapped reads were mapped to the human GRCh38.p13 reference genome or the *D. melanogaster* dmel r6 reference genome with STAR and the parameters: --runThreadN 16 --readFilesCommand gunzip -c –k --limitBAMsortRAM 20000000000 --outFilterMultimapNmax 1 --outFilterScoreMin 10 --outSAMtype BAM SortedByCoordinate. SAMtools was used to retain the properly paired and mapped reads (-F 4). SAMtools view with the –s option was used to subsample all the bam files to the bam file containing the lowest number of reads and then to create strand-specific BAM files. The normalization factor was calculated as: (number of *D. melanogaster* reads) / (number of human reads + number of *D. melanogaster* reads). Spiked normalized bedGraph files were created with BEDtools genomecov and the options: -bg -5 –scale (1 / normalization factor). Bigwig files were generated from the BedGraph files with the bedGraphToBigWig tool.

**ChIP-seq analysis**

Adapters were trimmed with Cutadapt with the following options: --minimum-length 10 -q 15, 10 -j 16 -a AGATCGGAAGAGCACACGTCTGAACTCCAGTCA. Trimmed reads were mapped to the human GRCh38.p13 reference genome with STAR and the parameters: --runThreadN 16 --readFilesCommand gunzip -c -k –alignIntronMax 1 --limitBAMsortRAM 20000000000 --outSAMtype BAM SortedByCoordinate. SAMtools was used to retain the properly mapped reads and to remove PCR duplicates. Reads mapping to the DAC Exclusion List Regions (accession: ENCSR636HFF) were removed with BEDtools. FPKM-normalized bigwig files were created with deepTools2 bamCoverage tool with the parameters -bs 10 -p max –normalizeUsing RPKM -e 200.

**ATAC-seq analysis**

Adapters were trimmed with Cutadapt with the following options: --minimum-length 10 -q 15, 10 -j 16 -A GATCGTCGGACTGTAGAACTCTGAAC -a AGATCGGAAGAGCACACGTCTGAACTCCAGTCAC. Trimmed reads were mapped to the human GRCh38.p13 reference sequence with STAR and the parameters: --runThreadN 16 --readFilesCommand gunzip -c -k –alignIntronMax 1 --limitBAMsortRAM 20000000000 --outSAMtype BAM SortedByCoordinate. SAMtools was used to retain the properly mapped reads and to remove PCR duplicates. Reads mapping to the DAC Exclusion List Regions (accession: ENCSR636HFF) and to the mitochondrial genome were removed with BEDtools. FPKM-normalized bigwig files were created with deepTools2 bamCoverage tool with the parameters -bs 10 -p max –normalizeUsing RPKM -e.

**TT-seq analysis**

Low-quality bases from 3' end of paired-end raw reads were trimmed using cutadapt 4.2. The reads were aligned to the human genome assembly GRCh38 release 103 with its annotation using STAR 2.7.5. The mapped reads with primary alignment and MAPQ $\geq$ 20 were retained and used for the downstream analysis. Strand-specific bigwig files were generated using deepTools bamCoverage 3.5.1. To estimate elongation rates, the transition points that correspond to the 5' end of the inhibition waves at 30-min NVP-2 treatment were called using proRate package in R 4.2.3 with BAM files for 0h and 0.5h NVP-2 treatment. We selected genes with >60kb long as we assume that the distance between TSS and the inhibition wave is ~60kb (~2 kb/min for 30 min). To improve the accuracy of transition point calling, we chose genes that showed >70% reduction in the read density at the genetic intervals (TSS+1kb to TSS+20kb) upon 30-min NVP-2 treatment. This eliminates genes with high background signal that result in poor calling of the transition points. Genes with reproducible calling of the transition points from 2 replicates were kept (n = 130) and used for the downstream analysis.

**Metagene Analysis**

Metagene profiles of genes scaled to the same length were then generated with Deeptools2 computeMatrix tool with the following parameters: -bs 10 -p max -m 4000 -b 2500 -a 2500. The plotting data were obtained with plotProfile –outFileNameData tool. Graphs representing the POINT-seq, PRO-seq, and ChIP-seq (IP / Input) signals were then created with GraphPad Prism 10.1. Metagene profiles are shown as the average of two biological replicates.

**Reads quantification**

Reads quantification over genomic regions (gene body: TSS to TES, termination region: TES to TES + 2.5kb, replication regions) was performed on strand-specific bam files with BEDtools multicov and the options -s -split. The reads quantifications were then normalized for library size (either to 100 million mapped paired-end reads or the spike-in normalization factor, as indicated on the figures) and for the region's length. TSS and TES correspond to the 5' and 3'ends of the most expressed transcript, respectively.

**Termination index**

The transcription termination index (TI) is defined as: TI = log2((([TES, TES + 2,500]$_{\text{read counts}}$ * normalization factor) / 2,500) / (([TSS, TES]$_{\text{read counts}}$ * normalization factor) / (Gene body size (TES – TSS))). The normalization factor is defined in Reads quantification (see above).

**Splicing efficiency**

The splicing efficiency on POINT-seq was calculated as in previous study[17] by first parsing each bam file to obtain the list of spliced and unspliced reads with the awk command (awk '/^@/ || $6 ~ /N/' for spliced reads and awk '/^@/ || $6 !~ /N/' for unspliced reads). The splicing efficiency was then calculated as the number of spliced reads over total reads with BEDtools multicov and the options –s –split.

**Data visualization**

Boxplots and violin plots, representing min to max with first quartile, median, and third quartile values were made with GraphPad Prism 10.1.

**P-values and significance tests**

P-values were computed with Wilcoxon rank sum test or a Wilcoxon signed-rank test. Statistical tests were performed in GraphPad Prism 10.1.

**Proteomics analysis**

Database searching: All MS/MS samples were analyzed using PEAKS XPro Studio (Bioinformatics Solutions, Waterloo, ON Canada; version 10.6 (2020-12-21)). PEAKS XPro Studio was set up to search the HuALLEGFR database (15,000 entries) assuming the digestion enzyme trypsin and three max missed cleavages. PEAKS Studio was searched with a fragment ion mass tolerance of 0.020 Da, a parent tolerance of 15 PPM.

Criteria for protein identification: Scaffold (version Scaffold_5.3.0, Proteome Software Inc., Portland, OR) was used to validate MS/MS based peptide and protein identifications. Peptide identifications were accepted if they could be established at greater than 50.0% probability by the Peptide Prophet algorithm[18] with Scaffold delta-mass correction. Protein identifications were accepted if they could be established at greater than 50.0% probability and contained at least 1 identified peptide. Protein probabilities were assigned by the Protein Prophet algorithm[19]. Proteins that contained similar peptides and could not be differentiated based on MS/MS analysis alone were grouped to satisfy the principles of parsimony.

Quantitative analysis: Scaffold was used to calculate the fold change and to perform t-test with Benjamini-Hochberg multiple test correction using as settings the quantitative method “Total Spectra” and “Use Normalization”. An adjusted p-value $< 0.05$ and a fold change $\geq |2|$ was considered significantly affected by the loss of NELF-C.

### SUPPLEMENTARY FIGURE LEGENDS

**Figure S1. Expression of *NELFCD* transcripts is highly up-regulated in colorectal cancer cells (Related to Figure 1).**

(A) Volcano plot of the log2 fold change of Tumors (T) vs Normal tissues (N) on 835 cell cycle-related genes in COAD. *CDKN1A*, *CDKN1B*, and *CDKN1C* genes are indicated in red.

(B) Comparative proteomic analysis of NELF subunits and SPT4 in human primary colon cancers and its adjacent tissues.

(C) Log2 of the RNA expression levels of *NELFCD*, *SUPT4H1*, *CDKN1A*, and *CDKN1C* genes in N and T of the indicated tissues and tumors are compared.

**Figure S2. Nuclear NELF complex controls the cell cycle (Related to Figure 2).**

(A) Western blot of NELF-C-AID DLD-1 whole cell extract (WCE) using the indicated antibodies. Treatment time (h) of IAA is also indicated.

(B) Western blot of SPT4-AID DLD-1 WCE using the indicated antibodies. Treatment time (h) of IAA is also indicated.

(C) MA plot of differential expression analysis of nuclear pA+ RNA-seq in NELF-C-AID DLD-1 cells (4h vs 0h IAA).

(D) Fold increase of nuclear pA+ RNA-seq on RDH genes in NELF-C-AID DLD-1 cells (0, 4, 12, 24h IAA).

(E) Western blot of NELF-C-AID DLD-1 cell nuclear fraction using the indicated antibodies. Treatment time (h) of IAA is also indicated.

**Figure S3. Acute NELF depletion fails to terminate Pol II transcription (Related to Figure 3).**

(A) Western blot of NELF-E-AID DLD-1 WCE and nuclear fraction using the indicated antibodies. Loading volume (x2 or x1) and treatment time (h) of IAA is also indicated. Tubulin protein was checked as a cytoplasmic marker.

(B) Western blot of NELF-C-AID DLD-1 WCE (4h DMSO, PlaB, IAA, PlaB+IAA) using the indicated antibodies.

(C) Example view of POINT-seq on *FAM120B* genes in NELF-C-AID DLD-1 cells (4h DMSO, PlaB, IAA, PlaB+IAA).

(D) Quantification of splicing fraction of POINT-seq in NELF-C-AID DLD-1 cells (4h DMSO, PlaB, IAA, PlaB+IAA).

(E) Example view of PRO-seq on *RPS23* genes in NELF-C-AID DLD-1 cells (0, 4, and 24h IAA). Previously published PRO-seq data[2] was re-analyzed. Termination defect was observed at 4h IAA.

(F) Boxplots of log2(TI) from PRO-seq in NELF-C-AID DLD-1 cells (0, 1, 4, and 24h IAA). Published PRO-seq data[2] was re-analyzed.

(G) Example views of TT-seq on *RPS23* and *HELLS* genes in NELF-C-AID DLD-1 cells treated for 4h with DMSO or IAA.

(H) Box plots of log2(TI) derived from TT-seq in NELF-C-AID DLD-1 cells treated for 4h with DMSO or IAA. Data for 2 biological replicates (R1 and R2) are shown. Statistical test: Wilcoxon rank sum test.

**Figure S4. NELF controls transcription termination independently of promoter proximal Pol II pausing or 3' poly(A) sites (Related to Figure 4).**

(A) Box plots of log2(TI) in POINT-seq in NELF-C-AID DLD-1 cells (4h vs 0h IAA). Genes were classified to UncTI, IncTI, and DecTI categories by TI.

(B) Metagene analysis of previously published NELF-C ChIP-seq[2] for UncTI and IncTI genes in NELF-C-AID DLD-1 cells.

(C) Metagene analysis of ATAC-seq for UncTI and IncTI genes in NELF-C-AID DLD-1 cells (0, 4, and 24h IAA).

(D) Quantification of ATAC-seq signals at TSS for UncTI and IncTI genes in NELF-C-AID DLD-1 cells (0, 4, and 24h IAA).

(E) Nucleotide distribution of adjacent sequences to PAS in UncTI and IncTI genes.

(F) Nucleotide composition of adjacent sequences to PAS in UncTI and IncTI genes.

(G) Profiles of TT-seq mean signal in NELF-C-AID DLD-1 cells treated for 4h with DMSO or IAA, followed by NVP-2 treatment for 0, 0.5, or 1h. Data for 2 biological replicates with ≥300 kb genes are shown.

**Figure S5. NELF loss alters 3'end processing of pc genes. (Related to Figure 5).**

(A) Heatmap analysis of nuclear pA+ RNA-seq in NELF-C-AID DLD-1 cells (4, 12, and 24h IAA). Fold change to 0h IAA is displayed in the indicated color.

(B) Fold change of RNA expression level of the indicated RDH genes in NELF-C-AID DLD-1 cells (0, 4, 12, and 24h IAA). *GAPDH* gene expression was not changed.

(C) View of POINT-seq of RDH gene cluster of NELF-C-AID DLD-1 cells (4h vs 0h IAA). Both (+) and (-) strands are shown.

(D) Metagene analysis of POINT-seq on RDH genes of NELF-C-AID DLD-1 cells (4h vs 0h IAA).

(E) Example view of nuclear pA+ RNA-seq and SIRV-normalized POINT-seq on *RPS23 and CTPS1* genes in NELF-C-AID DLD-1 cells (0, 4,12, and 24h IAA). PASs are indicated by arrows.

**Figure S6. Pol II transcription invades RI zones in the absence of NELF (Related to Figure 6).**

(A) Example view of SIRV-normalized POINT-seq on *FOCAD* gene adjacent to RI zone in NELF-C-AID DLD-1 cells (4h DMSO and IAA). Normalized POINT-seq signals are cut-off at 2000. RI zone is highlighted in gray. RI score: 22.7711. Both (+) and (-) strands are shown.

(B) Metagene analysis of POINT-seq on RI zones -/+2.5kb in NELF-C-AID DLD-1 cells (0, 4, 12, 24h IAA).

(C) Box plots of normalized POINT-seq signals in RI zones in NELF-E-AID DLD-1 cells (0h and 4h IAA). Two replicates are shown.

(D) Box plots of normalized POINT-seq signals in RI zones in SPT4-AID DLD-1 cells (0h and 3h IAA). Two replicates are shown.

(E) Box plots of normalized POINT-seq signals in RI zones in NELF-C-AID DLD-1 cells (4h DMSO and PlaB). Two replicates are shown.

(F) Example view of SIRV-normalized POINT-seq on *SPOPL* gene adjacent to P-NA zone in NELF-C-AID DLD-1 cells (4h DMSO and IAA). RI zone is highlighted in gray. RI score: 42.8401. Both (+) and (-) strands are shown.

(G) Box plots of TIs in POINT-seq in NELF-C-AID DLD-1 cells (4h vs 0h IAA). Genes in P-NA and P-A (all, all without RDH, and RDH genes) zones were analyzed.

**Figure S7. Rapid depletion of NELF-C protein causes global loss of DNA replication initiation factors on chromatin (Related to Figure 7).**

(A) Heatmap analysis of NELF proteins (0h and 4h IAA). Fold change to 0h IAA is displayed in the indicated color.

(B) Heatmap analysis of CPA and Terminator proteins (0h and 4h IAA). Fold change to 0h IAA is displayed in the indicated color.

**Supplementary Table 1. Number of GTEx and TCGA samples used for the analysis shown in Figure 1**

**Supplementary Table 2. Results of the proteomics performed on chromatin fractions of DLD-1 NELF-C-AID cells treated with DMSO or IAA for 4h**

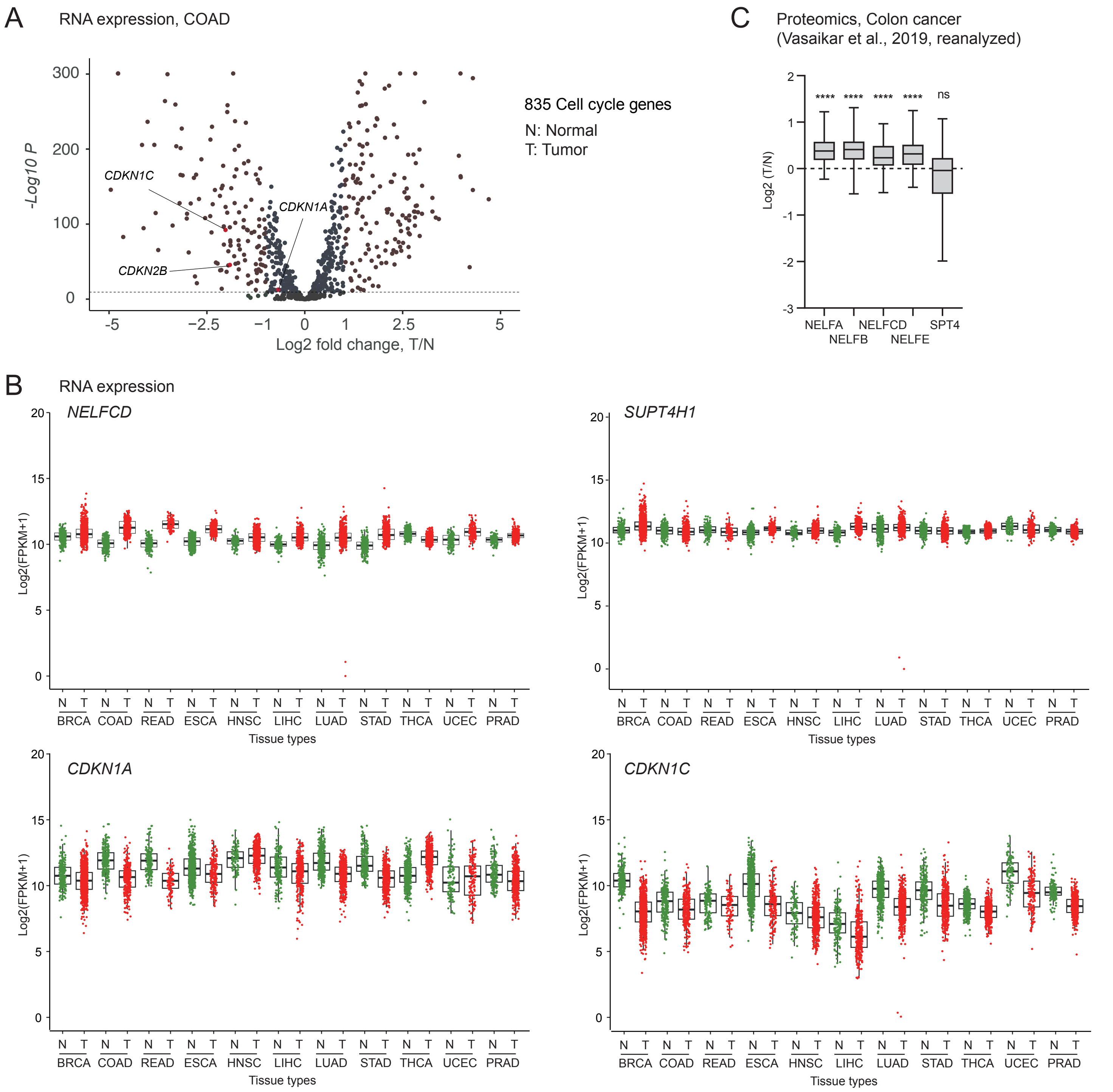
A
RNA expression, COAD
-Log10 P
Log2 fold change, T/N
CDKN1C
CDKN1A
CDKN2B
835 Cell cycle genes
N: Normal
T: Tumor
C
Proteomics, Colon cancer
(Vasaikar et al., 2019, reanalyzed)
Log2 (T/N)
**** **** **** **** ns
NELFA NELFB NELFCD NELFE SPT4
B
RNA expression
NELFCD
SUPT4H1
CDKN1A
CDKN1C
Log2(FPKM+1)
BRCA COAD READ ESCA HNSC LIHC LUAD STAD THCA UCEC PRAD
Tissue types

Figure S1

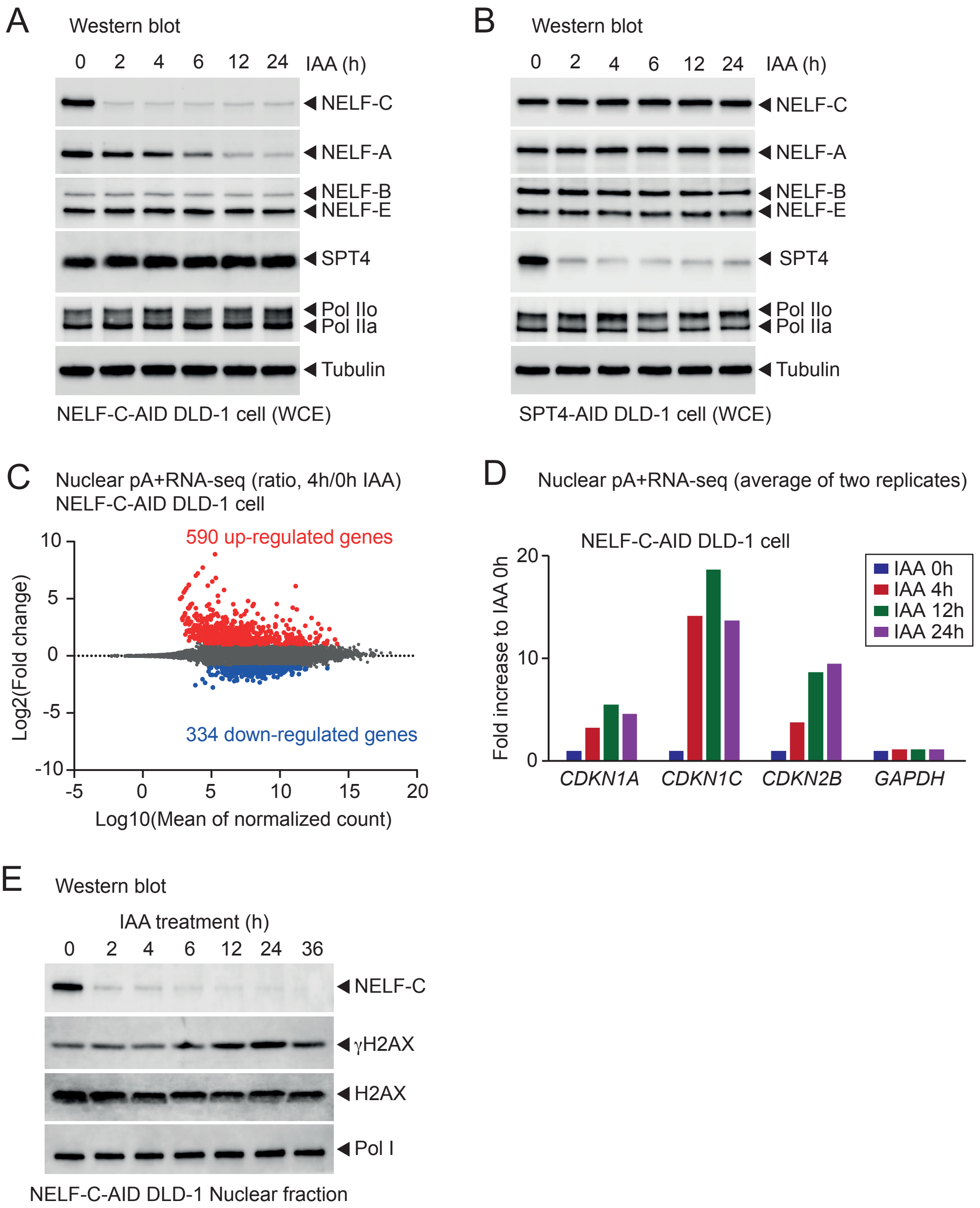
A
Western blot
0 2 4 6 12 24 IAA (h)
NELF-C
NELF-A
NELF-B
NELF-E
SPT4
Pol IIo
Pol IIa
Tubulin
NELF-C-AID DLD-1 cell (WCE)
B
Western blot
0 2 4 6 12 24 IAA (h)
NELF-C
NELF-A
NELF-B
NELF-E
SPT4
Pol IIo
Pol IIa
Tubulin
SPT4-AID DLD-1 cell (WCE)
C
Nuclear pA+RNA-seq (ratio, 4h/0h IAA)
NELF-C-AID DLD-1 cell
590 up-regulated genes
334 down-regulated genes
Log2(Fold change)
Log10(Mean of normalized count)
10
5
0
-5
-10
-5 0 5 10 15 20
D
Nuclear pA+RNA-seq (average of two replicates)
NELF-C-AID DLD-1 cell
Fold increase to IAA 0h
20
10
0
IAA 0h
IAA 4h
IAA 12h
IAA 24h
CDKN1A
CDKN1C
CDKN2B
GAPDH
E
Western blot
IAA treatment (h)
0 2 4 6 12 24 36
NELF-C
γH2AX
H2AX
Pol I
NELF-C-AID DLD-1 Nuclear fraction

Figure S2

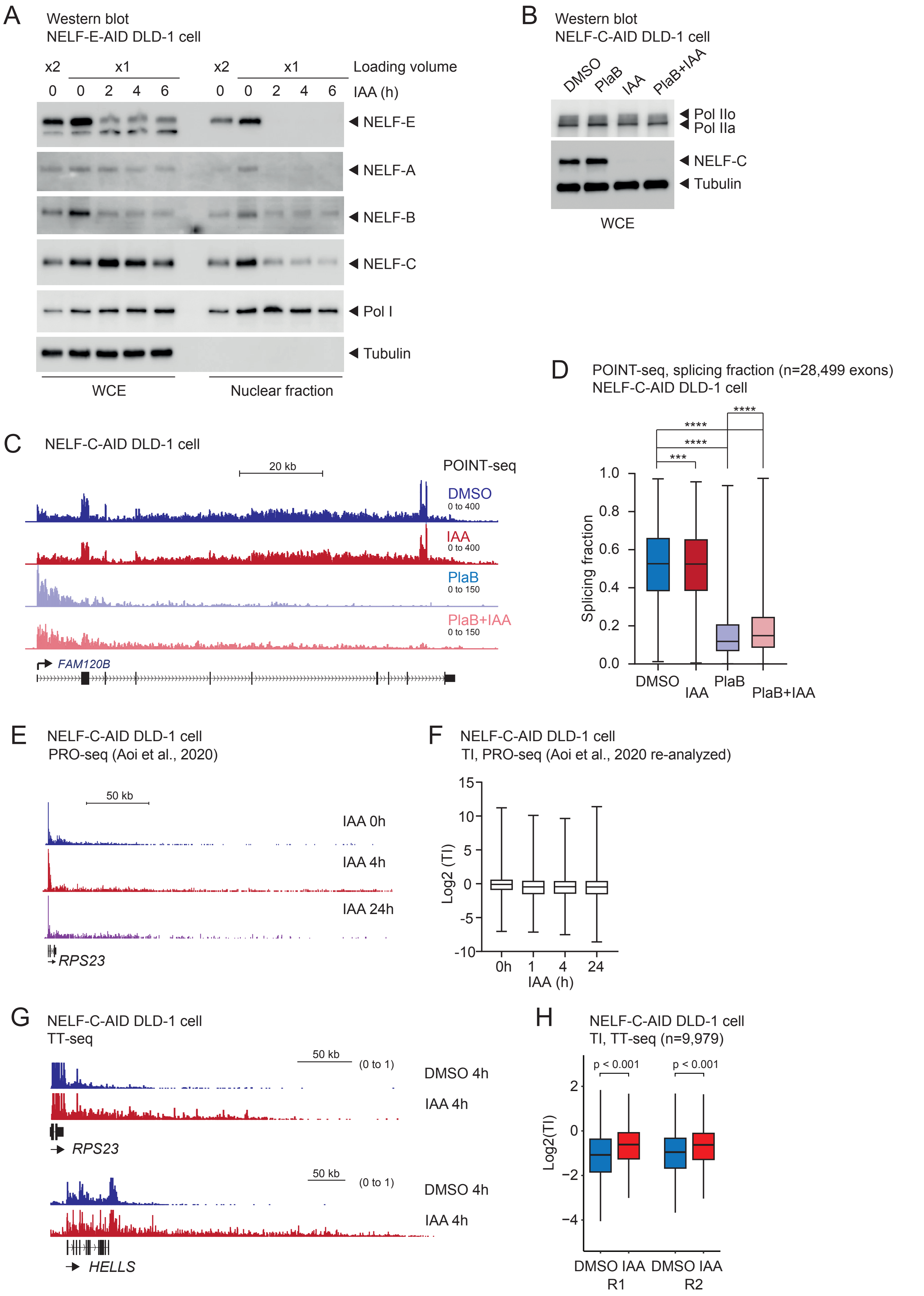

A
Western blot
NELF-E-AID DLD-1 cell
x2
x1
x2
x1
Loading volume
0 0 2 4 6 0 0 2 4 6
IAA (h)
NELF-E
NELF-A
NELF-B
NELF-C
Pol I
Tubulin
WCE
Nuclear fraction
B
Western blot
NELF-C-AID DLD-1 cell
DMSO
PlaB
IAA
PlaB+IAA
Pol IIo
Pol IIa
NELF-C
Tubulin
WCE
C
NELF-C-AID DLD-1 cell
20 kb
POINT-seq
DMSO
0 to 400
IAA
0 to 400
PlaB
0 to 150
PlaB+IAA
0 to 150
FAM120B
D
POINT-seq, splicing fraction (n=28,499 exons)
NELF-C-AID DLD-1 cell
****
****
****
***
Splicing fraction
1.0
0.8
0.6
0.4
0.2
0.0
DMSO
IAA
PlaB
PlaB+IAA
E
NELF-C-AID DLD-1 cell
PRO-seq (Aoi et al., 2020)
50 kb
IAA 0h
IAA 4h
IAA 24h
RPS23
F
NELF-C-AID DLD-1 cell
TI, PRO-seq (Aoi et al., 2020 re-analyzed)
Log2 (TI)
15
10
5
0
-5
-10
0h
1
4
24
IAA (h)
G
NELF-C-AID DLD-1 cell
TT-seq
50 kb
(0 to 1)
DMSO 4h
IAA 4h
RPS23
50 kb
(0 to 1)
DMSO 4h
IAA 4h
HELLS
H
NELF-C-AID DLD-1 cell
TI, TT-seq (n=9,979)
p < 0.001
p < 0.001
Log2(TI)
2
0
−2
−4
DMSO IAA
R1
DMSO IAA
R2


Figure S3

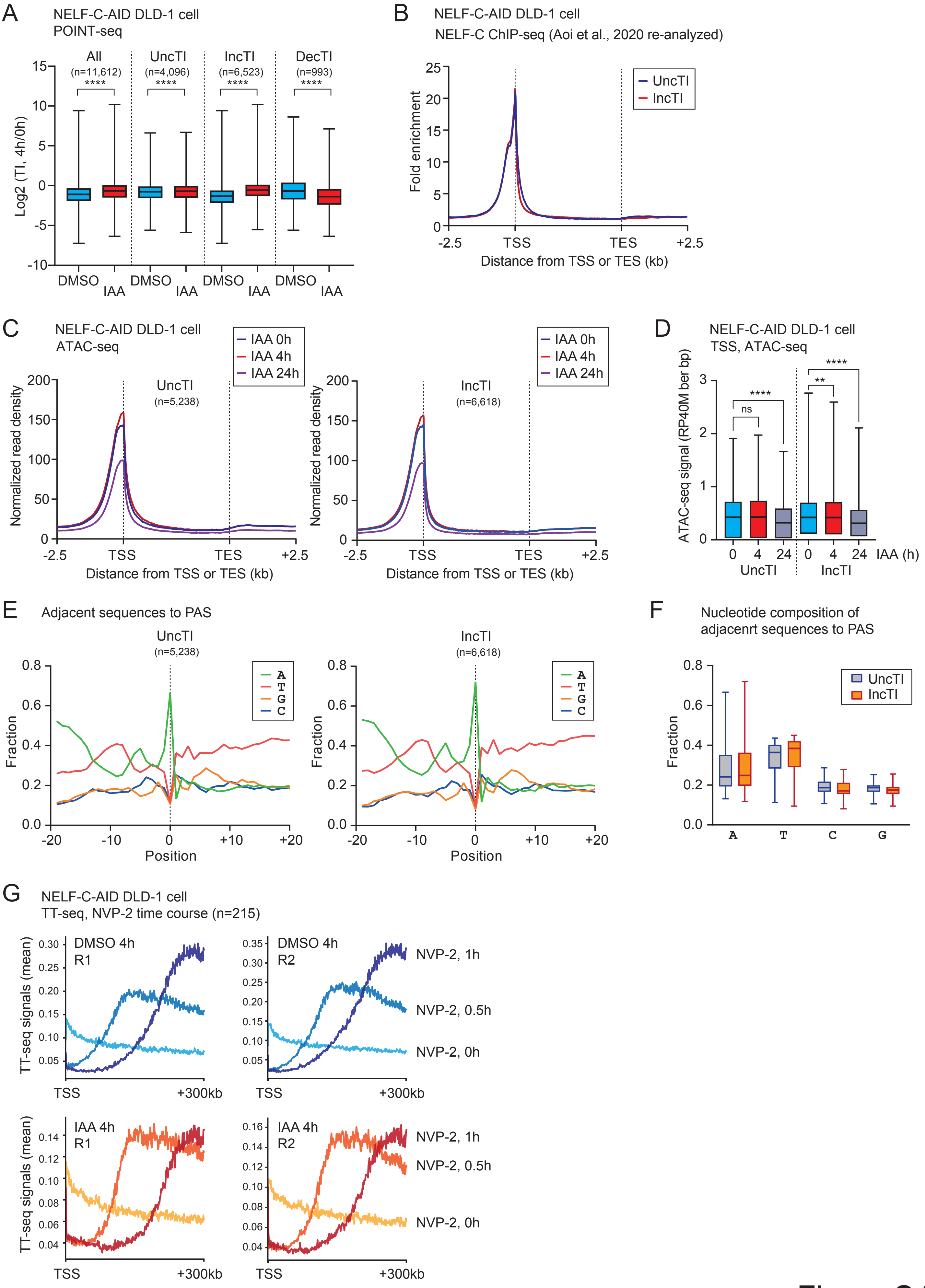
A
NELF-C-AID DLD-1 cell
POINT-seq
All
(n=11,612)
UncTI
(n=4,096)
IncTI
(n=6,523)
DecTI
(n=993)
****
Log2 (TI, 4h/0h)
DMSO
IAA
B
NELF-C-AID DLD-1 cell
NELF-C ChIP-seq (Aoi et al., 2020 re-analyzed)
UncTI
IncTI
Fold enrichment
-2.5
TSS
TES
+2.5
Distance from TSS or TES (kb)
C
NELF-C-AID DLD-1 cell
ATAC-seq
IAA 0h
IAA 4h
IAA 24h
UncTI
(n=5,238)
IncTI
(n=6,618)
Normalized read density
Distance from TSS or TES (kb)
D
NELF-C-AID DLD-1 cell
TSS, ATAC-seq
ATAC-seq signal (RP40M ber bp)
ns
**
****
IAA (h)
UncTI
IncTI
E
Adjacent sequences to PAS
UncTI
(n=5,238)
IncTI
(n=6,618)
A
T
G
C
Fraction
Position
F
Nucleotide composition of
adjacenrt sequences to PAS
UncTI
IncTI
Fraction
A
T
C
G
G
NELF-C-AID DLD-1 cell
TT-seq, NVP-2 time course (n=215)
DMSO 4h
R1
DMSO 4h
R2
IAA 4h
R1
IAA 4h
R2
NVP-2, 1h
NVP-2, 0.5h
NVP-2, 0h
TT-seq signals (mean)
TSS
+300kb

Figure S4

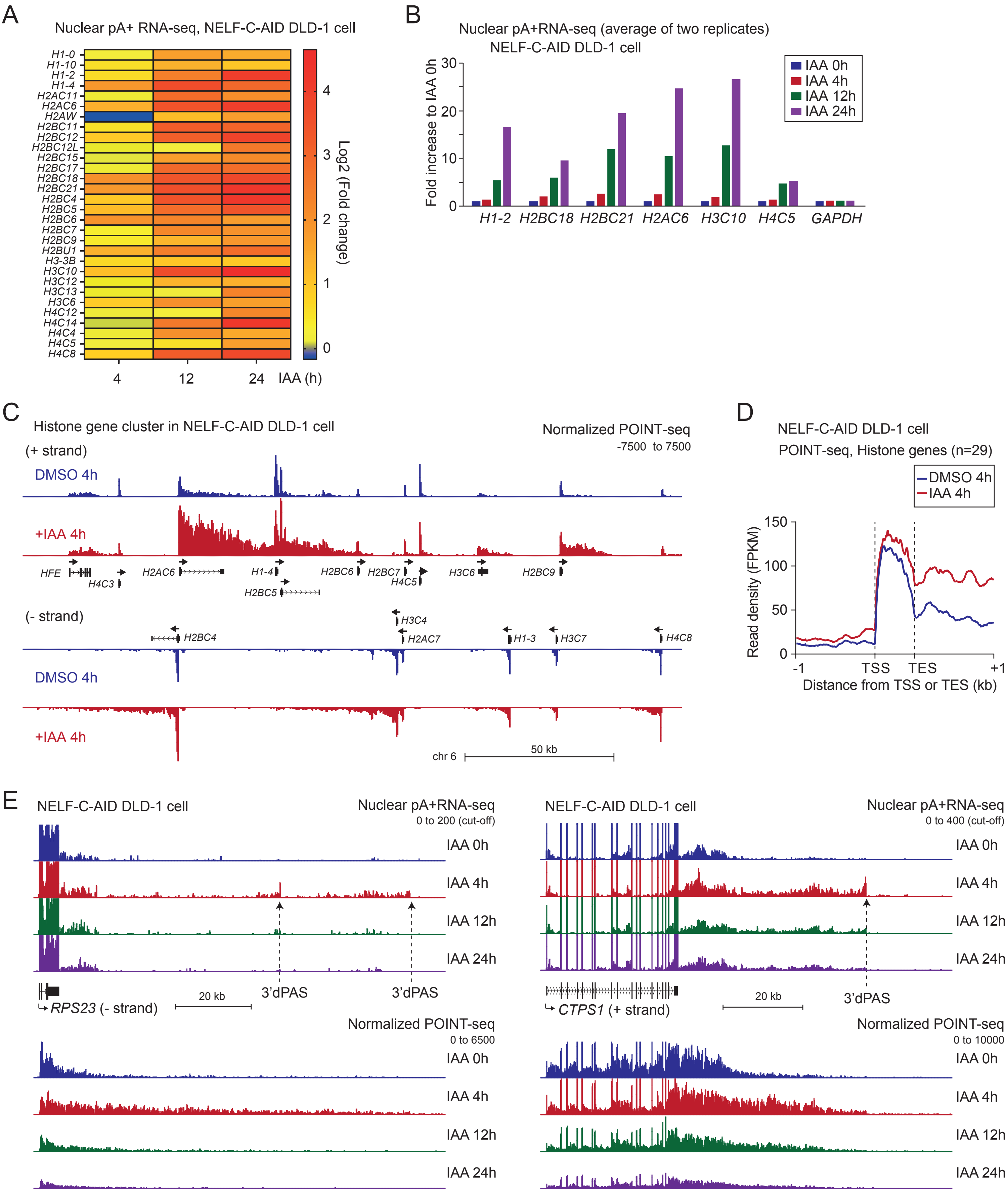
A
Nuclear pA+ RNA-seq, NELF-C-AID DLD-1 cell
H1-0
H1-10
H1-2
H1-4
H2AC11
H2AC6
H2AW
H2BC11
H2BC12
H2BC12L
H2BC15
H2BC17
H2BC18
H2BC21
H2BC4
H2BC5
H2BC6
H2BC7
H2BC9
H2BU1
H3-3B
H3C10
H3C12
H3C13
H3C6
H4C12
H4C14
H4C4
H4C5
H4C8
4
12
24
IAA (h)
Log2 (Fold change)
B
Nuclear pA+RNA-seq (average of two replicates)
NELF-C-AID DLD-1 cell
Fold increase to IAA 0h
IAA 0h
IAA 4h
IAA 12h
IAA 24h
H1-2
H2BC18
H2BC21
H2AC6
H3C10
H4C5
GAPDH
C
Histone gene cluster in NELF-C-AID DLD-1 cell
Normalized POINT-seq
-7500 to 7500
(+ strand)
DMSO 4h
+IAA 4h
HFE
H4C3
H2AC6
H1-4
H2BC5
H2BC6
H2BC7
H4C5
H3C6
H2BC9
(- strand)
H2BC4
H3C4
H2AC7
H1-3
H3C7
H4C8
DMSO 4h
+IAA 4h
chr 6
50 kb
D
NELF-C-AID DLD-1 cell
POINT-seq, Histone genes (n=29)
DMSO 4h
IAA 4h
Read density (FPKM)
-1
TSS
TES
+1
Distance from TSS or TES (kb)
E
NELF-C-AID DLD-1 cell
Nuclear pA+RNA-seq
0 to 200 (cut-off)
IAA 0h
IAA 4h
IAA 12h
IAA 24h
3'dPAS
3'dPAS
RPS23 (- strand)
20 kb
Normalized POINT-seq
0 to 6500
IAA 0h
IAA 4h
IAA 12h
IAA 24h
NELF-C-AID DLD-1 cell
Nuclear pA+RNA-seq
0 to 400 (cut-off)
IAA 0h
IAA 4h
IAA 12h
IAA 24h
CTPS1 (+ strand)
20 kb
3'dPAS
Normalized POINT-seq
0 to 10000
IAA 0h
IAA 4h
IAA 12h
IAA 24h

Figure S5

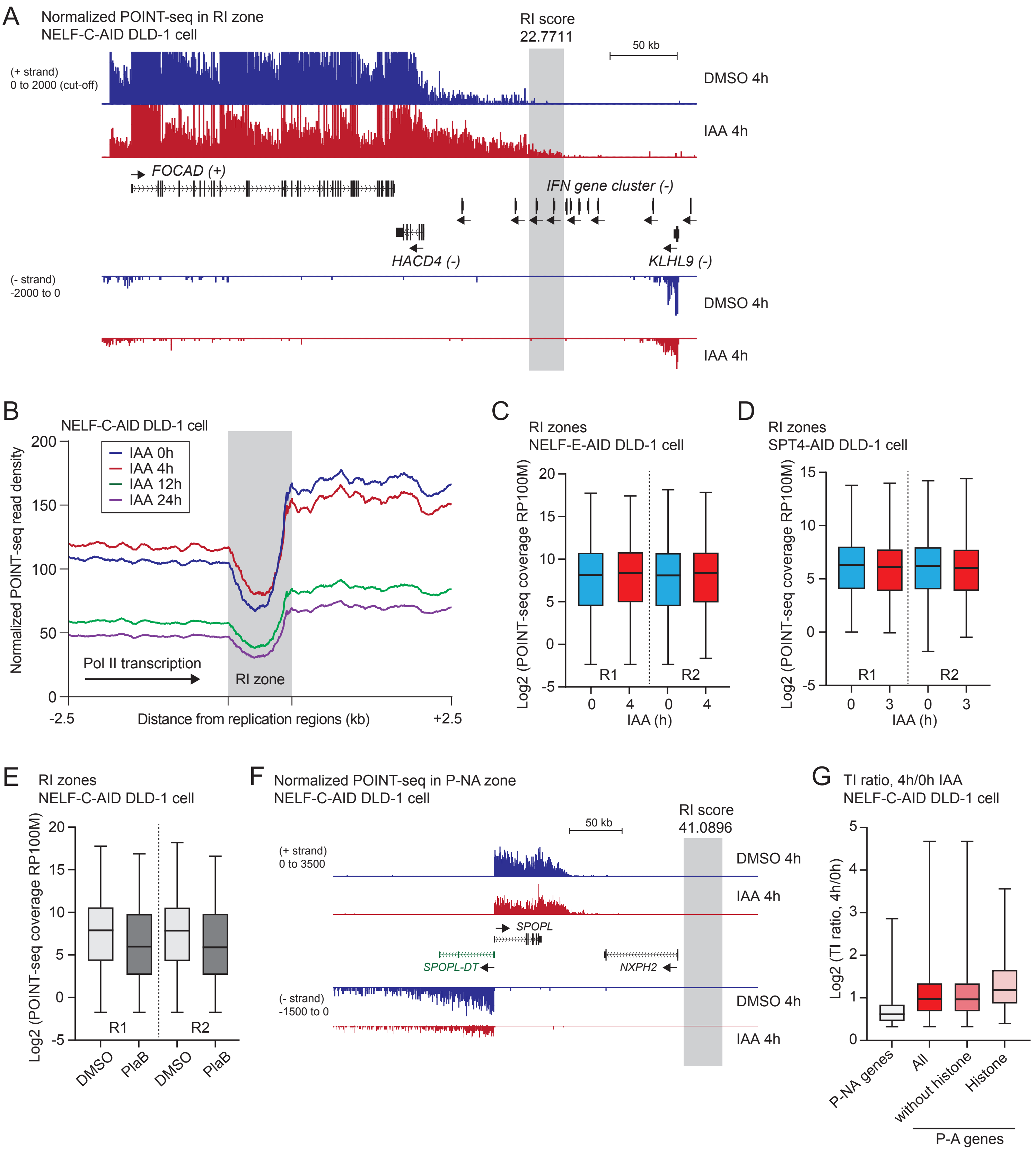

A
Normalized POINT-seq in RI zone
NELF-C-AID DLD-1 cell
RI score
22.7711
50 kb
(+ strand)
0 to 2000 (cut-off)
DMSO 4h
IAA 4h
FOCAD (+)
IFN gene cluster (-)
HACD4 (-)
KLHL9 (-)
(- strand)
-2000 to 0
DMSO 4h
IAA 4h
B
NELF-C-AID DLD-1 cell
IAA 0h
IAA 4h
IAA 12h
IAA 24h
Normalized POINT-seq read density
Pol II transcription
RI zone
Distance from replication regions (kb)
-2.5
+2.5
C
RI zones
NELF-E-AID DLD-1 cell
Log2 (POINT-seq coverage RP100M)
R1
R2
IAA (h)
D
RI zones
SPT4-AID DLD-1 cell
Log2 (POINT-seq coverage RP100M)
R1
R2
IAA (h)
E
RI zones
NELF-C-AID DLD-1 cell
Log2 (POINT-seq coverage RP100M)
R1
R2
DMSO
PlaB
DMSO
PlaB
F
Normalized POINT-seq in P-NA zone
NELF-C-AID DLD-1 cell
50 kb
RI score
41.0896
(+ strand)
0 to 3500
DMSO 4h
IAA 4h
SPOPL
SPOPL-DT
NXPH2
(- strand)
-1500 to 0
DMSO 4h
IAA 4h
G
TI ratio, 4h/0h IAA
NELF-C-AID DLD-1 cell
Log2 (TI ratio, 4h/0h)
P-NA genes
All
without histone
Histone
P-A genes


Figure S6

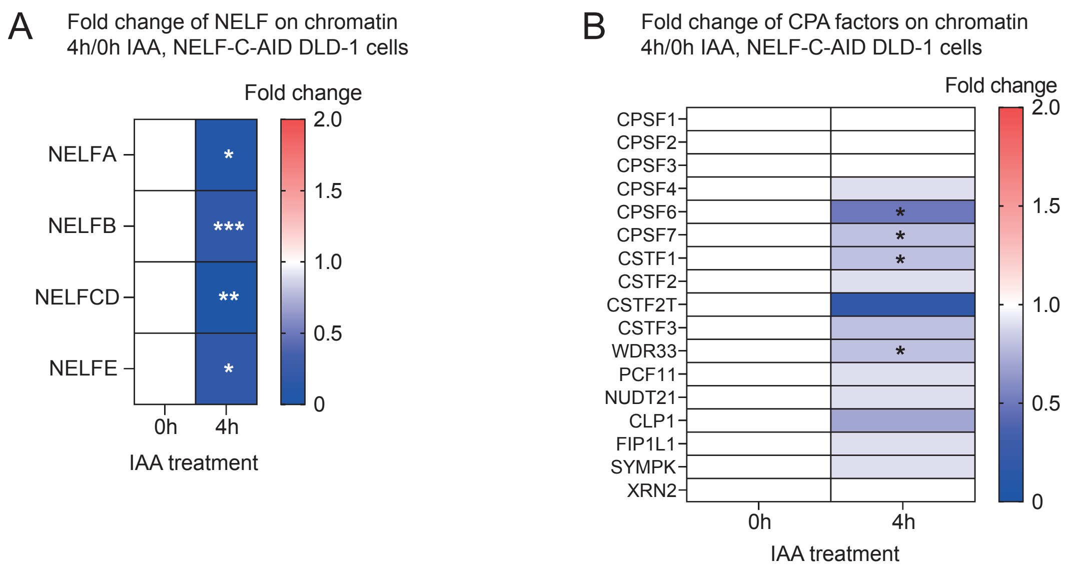
A
Fold change of NELF on chromatin
4h/0h IAA, NELF-C-AID DLD-1 cells
Fold change
NELFA
NELFB
NELFCD
NELFE
*
***
**
*
2.0
1.5
1.0
0.5
0
0h
4h
IAA treatment
B
Fold change of CPA factors on chromatin
4h/0h IAA, NELF-C-AID DLD-1 cells
Fold change
CPSF1
CPSF2
CPSF3
CPSF4
CPSF6
CPSF7
CSTF1
CSTF2
CSTF2T
CSTF3
WDR33
PCF11
NUDT21
CLP1
FIP1L1
SYMPK
XRN2
2.0
1.5
1.0
0.5
0
0h
4h
IAA treatment

Figure S7
